# Concerted epithelial and stromal changes during progression of Barrett’s Esophagus to invasive adenocarcinoma exposed by multi-scale, multi-omics analysis

**DOI:** 10.1101/2023.06.08.544265

**Authors:** Michael K. Strasser, David L. Gibbs, Philippe Gascard, Joanna Bons, John W. Hickey, Christian M. Schürch, Yuqi Tan, Sarah Black, Pauline Chu, Alican Ozkan, Nathan Basisty, Veena Sangwan, Jacob Rose, Samah Shah, Sophie Camilleri-Broet, Pierre-Oliver Fiset, Nicolas Bertos, Julie Berube, Haig Djambazian, Rui Li, Spyridon Oikonomopoulos, Daffolyn Rachael Fels-Elliott, Sarah Vernovsky, Elee Shimshoni, Deborah Collyar, Ann Russell, Ioannis Ragoussis, Matthew Stachler, James R. Goldenring, Stuart McDonald, Donald E. Ingber, Birgit Schilling, Garry P. Nolan, Thea D. Tlsty, Sui Huang, Lorenzo E. Ferri

**Author notes:** equal contribution (Michael K. Strasser, David L. Gibbs, Donald E. Ingber, Birgit Schilling, Garry P. Nolan, Thea D. Tlsty, Sui Huang, Lorenzo E. Ferri).

## Abstract

Esophageal adenocarcinoma arises from Barrett’s esophagus, a precancerous metaplastic replacement of squamous by columnar epithelium in response to chronic inflammation. Multi-omics profiling, integrating single-cell transcriptomics, extracellular matrix proteomics, tissue-mechanics and spatial proteomics of 64 samples from 12 patients’ paths of progression from squamous epithelium through metaplasia, dysplasia to adenocarcinoma, revealed shared and patient-specific progression characteristics. The classic metaplastic replacement of epithelial cells was paralleled by metaplastic changes in stromal cells, ECM and tissue stiffness. Strikingly, this change in tissue state at metaplasia was already accompanied by appearance of fibroblasts with characteristics of carcinoma-associated fibroblasts and of an NK cell-associated immunosuppressive microenvironment. Thus, Barrett’s esophagus progresses as a coordinated multi-component system, supporting treatment paradigms that go beyond targeting cancerous cells to incorporating stromal reprogramming.

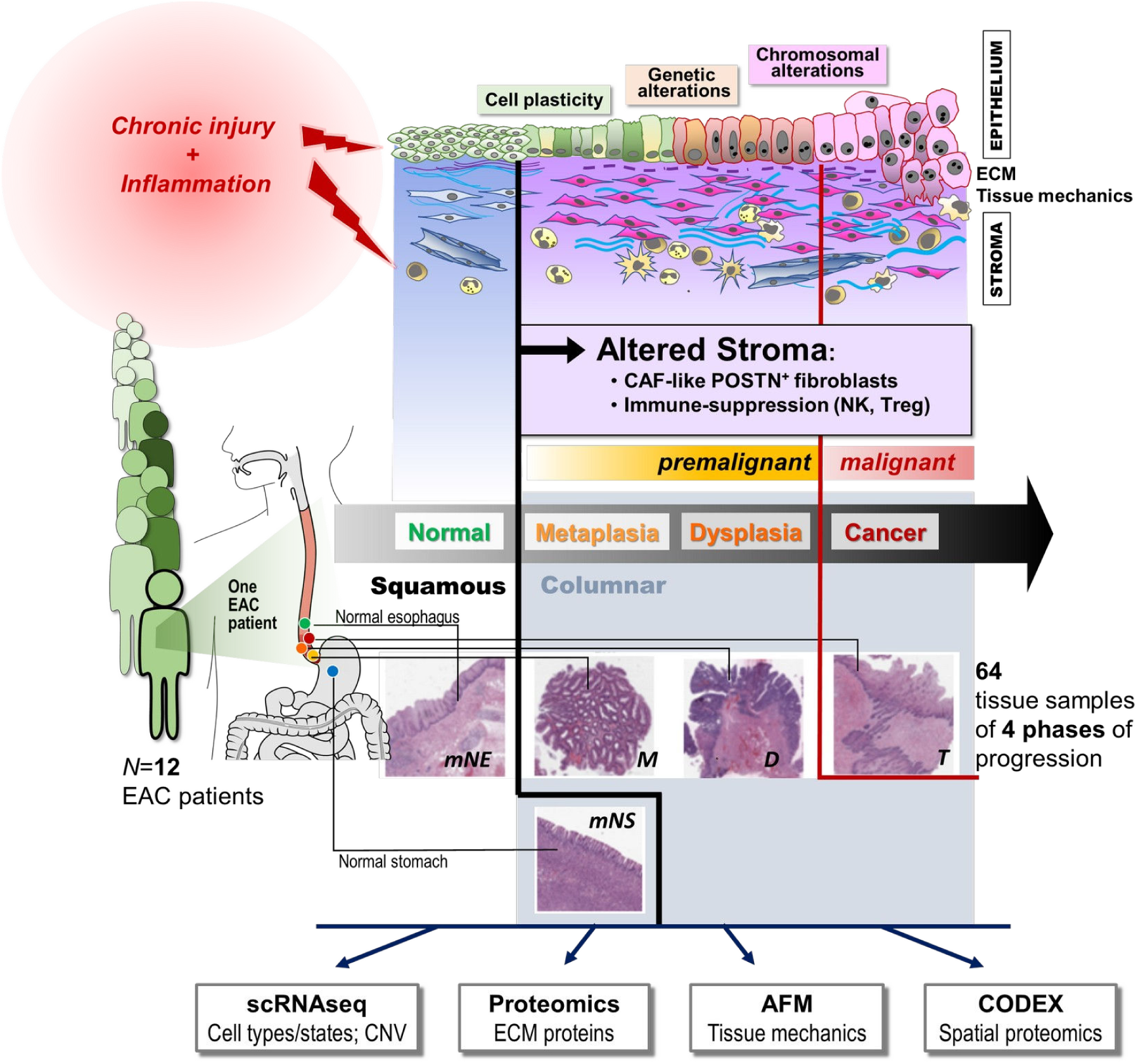
Graphical Abstract. To obtain a comprehensive picture of the coordinated changes in epithelial, stromal and immune compartments during development of Barrett’s-associated esophageal adenocarcinoma, patient-matched samples corresponding to various phases of disease progression were collected from 12 patients, each of which had at a given time point lesions at multiple stages progression (matched-normal, metaplasia, dysplasia, and carcinoma). Matched “normal” gastric tissues were also collected. These sample were analyzed by single cell RNA-sequencing (scRNAseq) for single-cell resolution transcriptomics and Copy Number Variant (CNV), by proteomics for extracellular matrix (ECM) proteins, by Atomic Force Microscopy (AFM for tissue stiffness and by CODEX spatial proteomics imaging The integrative multi-omics analysis exposed drastic alterations in cell type composition and shifts in cell states in all three compartments. A large subpopulation of fibroblasts absent in the normal esophagus and characteristic of dysplasia and adenocarcinoma sample, that based on markers would indeed be considered cancer associated fibroblasts (CAF), appeared already in the metaplastic phase. This fibroblast subpopulation had transcriptomes virtually indistinguishable with fibroblasts of the cancer free gastric epithelium in these patients

## INTRODUCTION

Esophageal cancer is the seventh most common cancer worldwide^1^ and can be divided into two major subtypes that constitute biologically distinct diseases: esophageal squamous cell carcinoma (ESCC), observed predominantly in the upper esophagus, and esophageal adenocarcinoma (EAC), typically located in the lower esophagus^2^. These subtypes differ in etiology, epidemiology and genetic characteristics. ESSC is the dominant type in much of Asia^3^ while EAC is the prevalent type in Western countries, with a rapidly increasing incidence of currently 5-10 cases/100,000/year^4, 5^.

Histologically, EAC frequently displays a glandular tissue structure and genomic alterations that are indistinguishable from those of the chromosomally unstable variant of proximal gastric cancer ^6^ and very distinct from the original squamous tissue structure typical of the esophagus. Most EACs typically develop from metaplasia, an adaptive change to chronic injury caused by gastro-esophageal reflux (GERD) of the lower mucosa of the esophagus known as Barrett’s esophagus (BE) ^7^. In this metaplastic adaptive response driven by chronic inflammation, native squamous epithelium of the normal esophagus is replaced by a columnar epithelium with gastric and/or intestinal characteristics and mucin secretion. The prevalence of BE in the general population in Western countries is estimated to be 2% and is typically only diagnosed incidentally in patients with symptomatic GERD who undergo endoscopy. Patients with BE have a 40-fold increased lifetime risk of developing EAC. However, only a small minority of BE cases (<1% per patient/year) progress to invasive carcinoma ^8^.

Recent genome-wide analyses have focused on the identification of molecular changes occurring in the esophagus epithelium during the emergence of BE to address the long-standing question of the cellular origin of the metaplastic cells^7^ and characterize early genomic, extrachromosomal ^9–13^ and transcriptomic alterations^14, 15^. However, increasing evidence suggest that stromal alterations due to chronic injury play a central role in development of metaplasia and progression to cancer^16–18^ . Therefore, it becomes paramount to investigate disease processes in the context of the whole tissue rather than focusing only on one of its components, i.e., the epithelium. To this end, we performed a systematic multi-omics analysis to characterize, in depth, the concomitant changes in epithelial, immune, and stromal cell landscape, as well as changes in composition of the extracellular matrix and tissue mechanics, during progression to cancer.

Chronic inflammation due to recurrent tissue injury can perturb tissue homeostasis and thereby contribute to tumorigenesis^19^ . Failure to restore the original tissue structure perpetuates an aberrant regenerative response in which the parenchymal cells do not return to a normal, stably-differentiated state. The tissue may enter a metaplastic state that constitutes an adaptive response. In the case of BE-associated EAC, the esophageal mucosa is exposed to prolonged bile salts and acid reflux, dietary irritants, alcohol, and smoking and the squamous epithelia of the lower esophagus is replaced by columnar gastro-intestinal epithelia. For poorly understood reasons, metaplasia, inflammation and stromal reorganization also place the tissues at a higher risk for malignancy. We have examined how this perturbed homeostasis may drive tumorigenesis. The multifocal nature of BE-associated EAC, repeated endoscopic surveillance and surgical resection in the absence of (neo)adjuvant therapy affords an unparalleled opportunity to obtain a unique cohort of patient-matched tissue samples corresponding to all clinical histological/diagnostic stages (hereafter, considered the biological ‘*phase’* of progression) that can be captured at a single time point in a given patient.

Our simultaneous multi-omics analysis of parenchyma and stroma at the various phases of progression of BE to EAC revealed a coordinated program of non-genetic plasticity that included the epithelium, the extracellular matrix (ECM) and stroma. This change in multi-component tissue identity at the metaplastic phase was accompanied by acquisition of an NK-associated immunosuppressive tissue state and the appearance of fibroblasts with characteristics previously associated with malignant stroma. The joint consideration of single-cell resolution and tissue level molecular profiles enabled us to link known and novel molecular markers of progression to their cellular origins and to shifts in cell-type composition. Together, this multi-scale integration offers a starting point for designing multi-pronged stromal reprogramming as a new therapeutic modality for reverting premalignancy or preventing malignant progression.

## RESULTS

### Patient Recruitment and Sample Characteristics

A unique cohort of patients with BE-associated esophageal EAC was prospectively recruited to participate in this study. A total of 64 fresh tissue samples from 12 newly diagnosed, treatment-naive patients with confirmed EAC were collected for multi-omics analysis (Figs. 1A & S1). The samples encompassed inflamed squamous esophagus (>5cm proximal to apical columnar-lined mucosa), suspected Barrett’s metaplasia (columnar-lined mucosa), suspected dysplasia (based on high resolution white light and narrow band imaging on endoscopy), and malignant tumor. In the majority of cases, we also collected columnar stomach (gastric cardia) tissue. Samples were collected at the time of endoscopy (5/12) or surgical resection (7/12). Histological confirmation of suspected tissue diagnosis/stage (phase of progression, see below) was provided by independent analysis by two expert pathologists and the samples were labeled accordingly (Methods). Patient information, sample description, and histological diagnosis are provided in Table S1.

**Figure 1.**
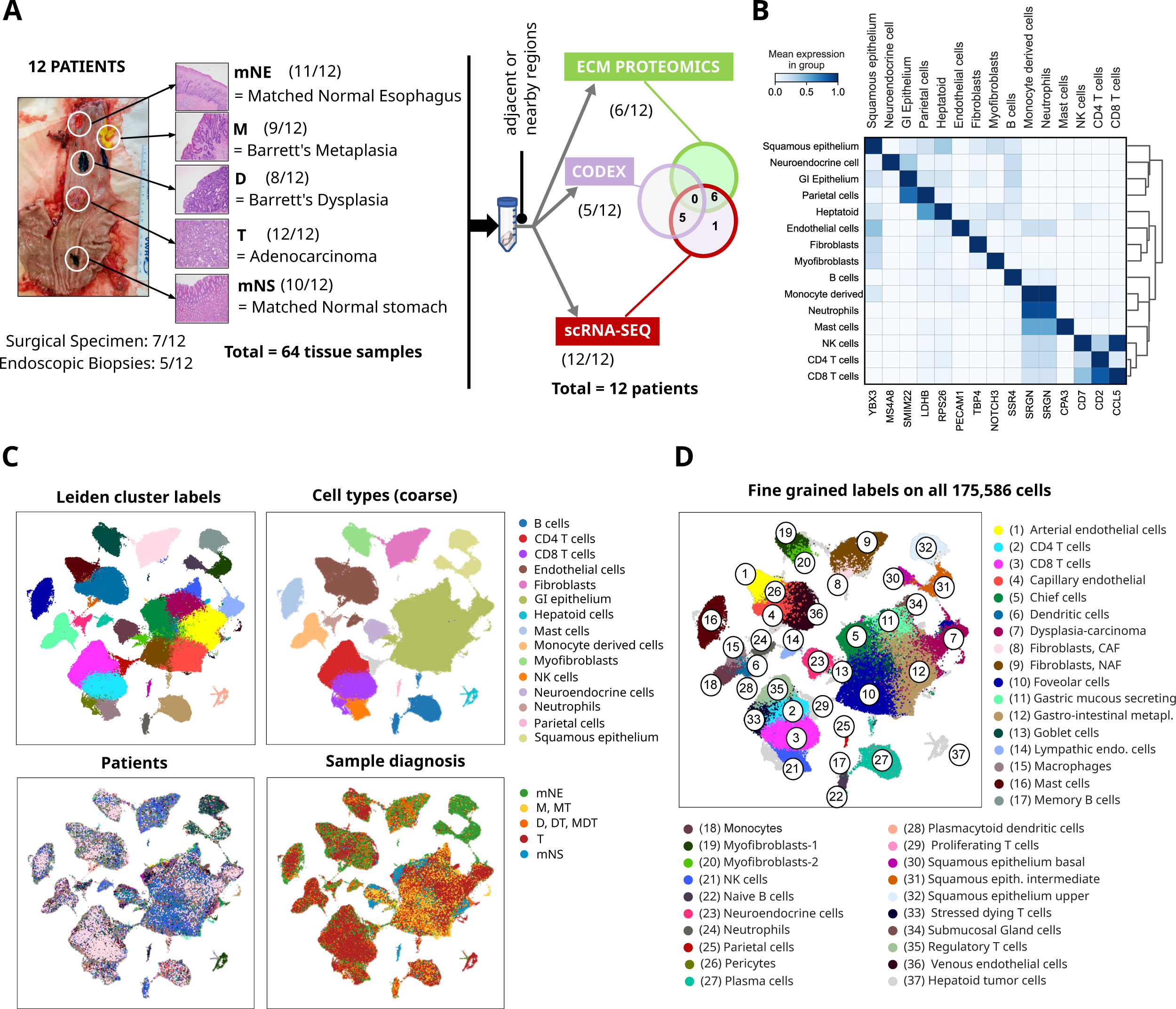
Integrated multiomics analysis of the progression of Barrett’s Esophagus (BE) to esophageal adenocarcinoma (EA). **(A)** Example of a surgical resection showing locations of samples representing the phases of disease progression and use in multiomic-analyses. **(B)** Gene markers (transcripts) used for classification of coarse-grained cell types from scRNAseq data. **(C)** UMAP visualization of the entirety of cells from all samples analyzed by scRNAseq color-labelled for clusters (Leiden unsupervised clustering), the coarse-grained cell types, tissue diagnosis, and patient IDs (clockwise from top left) obtained from scRNAseq. **(D)** UMAP visualization of all analyzed cells from al samples obtained from scRNAseq, color-labelled for the fine-grained cell type.

### Global survey of cell types in progression from inflamed esophagus to EAC

Transcriptomic profiling of 64 samples collected from 12 patients with EAC by single-cell RNA sequencing (scRNA-seq) captured the transcriptomes of 175,586 individual cells (Methods). This dataset consisted of patient-matched samples encompassing, in various combinations, the multiple phases of BE progression defined by the clinical-histological stage/diagnosis: matched “Normal” Esophagus (*mNE*) (obtained from the inflamed squamous esophageal tissue adjacent to the lesion), Barrett’s Metaplasia (*M*), Dysplasia (*D*), Tumor (*T*), and matched “Normal” Stomach (*mNS*) (obtained from the columnar gastric tissue adjacent to the lesion). We also collected samples diagnosed as mixed histology, i.e., with more than one diagnosis as frequently observed in this multifocal disease (Fig. S1B). For example, specimen *M*/*D* for patient E17 contained both metaplasia and dysplasia.

After preprocessing, dimension reduction and clustering (Methods), cells clustered mainly by cell type, but some patient/sample-specific variations remained despite our strictly standardized procedure for processing live specimens (Fig. S2A). The Harmony algorithm^20^ was therefore applied to remove such batch effects (Fig. S2B). The unsupervised cluster analysis of cells produced 40 clusters across the two clinical labels, diagnostic stage (progression phase) and patient. Correlation with reference cell type transcriptomes from the Human Cell Landscape (HCL) database^21^ (Methods) provided “coarse-grained” cell type labels for the majority of cells. Hence, the 40 clusters could be assigned to 16 readily identifiable cell types. The largest group of cells comprised gastro-intestinal (GI) epithelial cells (78K cells), followed by endothelial cells (17.7K), CD8 T cells (17.1K), CD4 T cells (14.3K), and fibroblasts (9.5K) (Fig. 1B-C, Table S2). In the description of this study, marker genes are designated as positively expressed unless explicitly annotated as negative.

The coarse-grained cell type clusters were then split into “fine-grained” categories representing 37 cell subtype labels (Fig. 1D). For example, the gastro-intestinal (GI) epithelial cells marked by MUC5AC expression were further divided into “functional types” (subtypes) that included goblet cells, chief cells, and foveolar cells. Gastric mucus-producing cells were identified by strong expression of MUC6 (75% of cells). Chief cells were identified by expression of PGC and LIPF, goblet cells through expression of MUC2, TFF3 and SPINK4 and foveolar cells with MUC1, MUC5AC and TFF1^22^ . For metaplasia, cells expressing MUC2 and CDX2 without MUC5AC were termed ‘intestinal metaplasia’^23^ . The batch-corrected cluster analysis thus correctly prioritized cell subtype over patient and progression phase as the discriminatory cluster feature. Therefore, epithelial cells from nominal dysplasia and tumor samples (*D*, *T*) were intermixed with normal and metaplastic cells in each of the cell type clusters; but the *D*, *T* cells had on average the highest fraction of markers for cell division, consistent with the diagnostic cell labels (Fig. S2 C-D). The lists of gene markers for each cell (sub)type based on statistical differential expression analysis between the clusters are available in Table S3.

**Figure 2.**
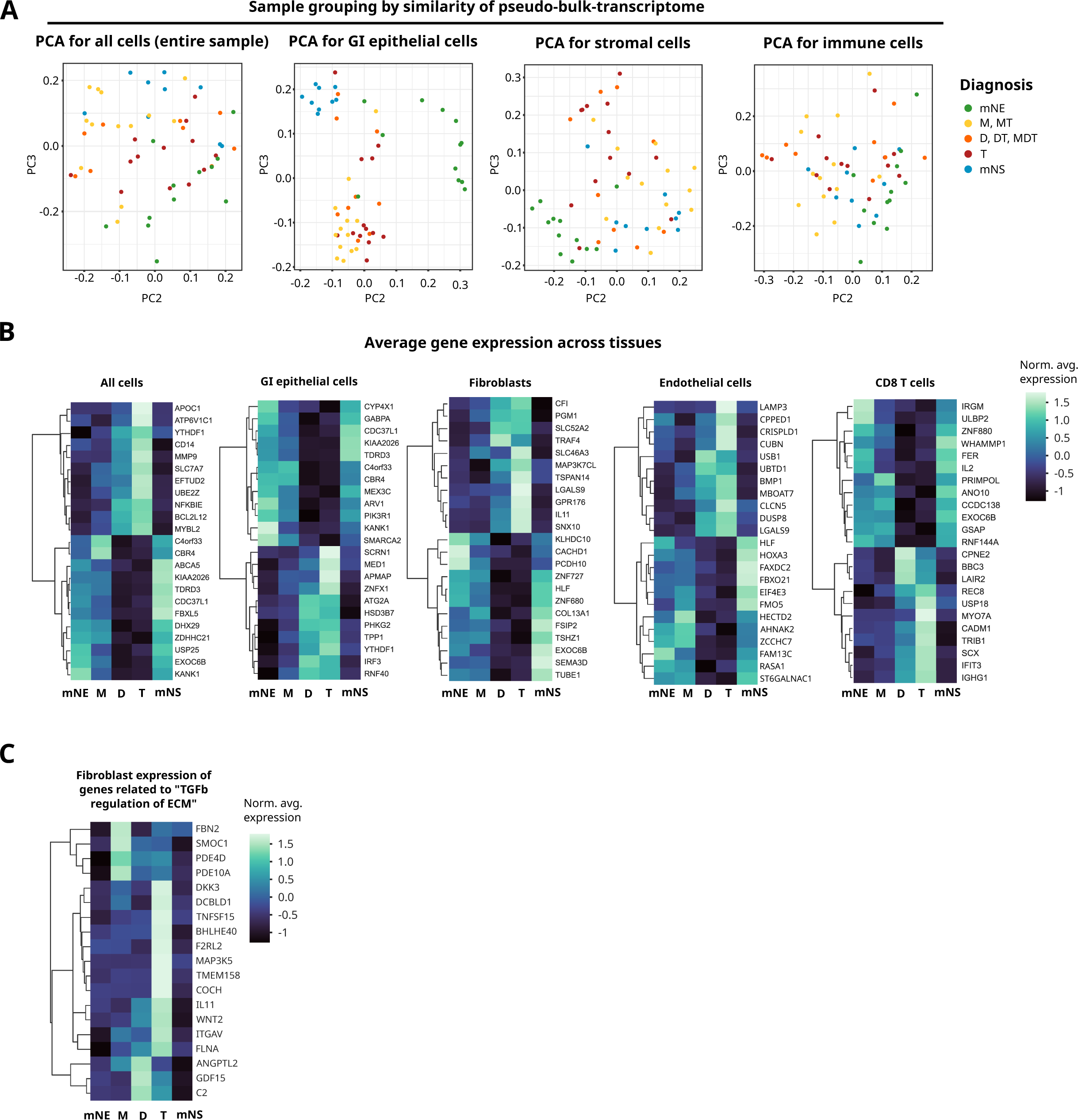
Overview of pseudo-bulk analysis. **(A)** Principal component analysis (PCA) displaying showing PC2 (x-axis) and PC3 (y-axis) of samples using pseudo-bulk transcriptome profiles for all cells (left panel), gastrointestinal (GI) epithelial cells (middle left panel), stromal cells (fibroblasts and endothelial cells) (middle right panel), and immune cells (T cells, B cells, Monocyte derived, Natural Killer (NK) cells, and dendritic cells (DC) (right panel)). Colors indicate the tissue diagnosis. **(B)** Heat maps showing changes in expression levels of selected genes (from left to right) for various pseudo-bulk schemes: all cells combined, gastro-intestinal (GI) epithelial cells, fibroblasts, endothelial cells and CD8 T cells at each phase of disease progression. **(C)** Heat map showing changes in expression levels of genes related to TGF-beta signaling in fibroblasts at each phase of disease progression.

### Transcriptome clusters during progression are predominantly determined by the epithelium

While single-cell transcriptomics allowed for resolving cell type heterogeneity, in order to map cell populations in their entirety to the transcriptome space and derive differential gene expression between progression phases, we computationally created “pseudo-bulk” transcriptomes^24^ selectively for the coarse-grained cell types of each sample (Methods). Principal Component Analyses (PCA) of pseudo-bulks transcriptomes that included all cell types (mimicking classical bulk RNAseq) failed to show diagnostic separation (Fig. 2A, left panel). However, PCA using only epithelial cell pseudo-bulk transcriptomes were able to group the various nominal histological diagnoses. Indeed, *mNE* samples, primarily inflamed squamous esophageal epithelium, were clearly separated from *M*, *D* and *T* along PC2, with the exception of sample E24A (Fig. 2A, middle left panel). By contrast, and as expected, the mNS samples were close to the cluster of *M*, *D* and *T* samples, and, in particular, overlapped with the *M* samples in PC2 dimension, consistent with the current paradigm on Barrett’s origin arising from proximal migration of gastric glandular mucosa^7^, but they were still separated in PC3 (Fig. 2A, middle left panel). Overall, the *M* and *D* samples clustered with the *T* samples (Fig. 2A).

The linear discrimination by PCA lessened, but still remained, when stromal (fibroblast/myofibroblasts/ endothelial) pseudo-bulk transcriptomes were used instead of epithelial cells (Fig. 2A, middle right panel). Importantly, there was an intermixing between fibroblast cells from *mN*S with those from *M*, *D*, and *T* samples, suggesting that fibroblast phenotype reflects an early shift from a squamous (esophageal) to a columnar cell context. Immune cell bulk transcriptomes did not discriminate the progression phases as clearly as stromal cells (Fig. 2A, right panel).

### Global Differential gene expression analysis reveal an immune suppressive stroma and novel dysplastic markers

To identify differentially expressed genes (DEGs) between the phases of progression, we used DESeq2^25^ and pseudo-bulk transcriptomes of the coarse-grained cell types (Table S2) (Methods). The largest group of cells, the GI epithelium, revealed the largest number of differentially expressed genes (6874 genes) between tissue diagnoses, followed by endothelial cells (1612 genes), fibroblasts (624 genes) and CD8 T cells (833 genes) (Figs. 2B & S3, Table S2).

In pathway enrichment analysis^26, 27^, DEGs of fibroblasts, B, Natural Killer (NK) and monocyte-derived cells displayed significant associations for particular pathways (Table S4). For example, fibroblast DEGs were strongly enriched for the functional annotation "TGF-beta regulation of extracellular matrix" (BioPlanet 2019, adj. p-value 6e-17) (Fig. 2C), supporting acquisition of a carcinoma-associated fibroblast (CAF) phenotype. Notably, genes characteristic of this pathway were under-expressed in both *mNE* and *mNS* samples compared to *M*, *D* and *T* samples (Table S4). Supporting pathway analysis, DEG in fibroblasts (increased SMOC1, FBN2, PDE4D and PDE10A in *M* samples; increased GDF15, ANGPTL2, C2 in *D* samples; and increased MAP3K5, TMEM158, COCH, TNFSF15, F2RL2, DKK3, BHLHE40, DCBLD1, ITGAV, IL11, WNT2, and FLNA in *T* samples) jointly point to a TGF-beta driven immunosuppressive and fibrotic program.

Although the goal of this survey was not to weigh in on evidence for the various hypotheses on the cell of origin of BE^28, 29^ our scRNAseq analysis confirmed the presence in our *M* phase samples of previously noted “mixed” phenotype (multi-lineage) epithelial cells in conjunction with this question (Fig. S3B-D), including transitional (squamous/columnar) basal progenitor cells (KRT5+/KRT7+)^7, 30^ as well as the gastric/intestinal mixed-phenotype cells^15^.

### Cell type composition changes during progression to EAC reflect a shift in tissue identity from esophagus to stomach

DEGs between phases can result either from changes in expression of genes in individual cells or from a shift in relative abundance of cell number (cell type composition change). Single cell-resolution transcriptomes permit comparison of samples at the level of the relative abundance of cell types instead of molecular profiles. To evaluate the discriminatory power of this higher-level feature, the similarities of the fine-grained cell type proportions were used to clusters the samples (Fig. S4A). Unsupervised hierarchical clustering grouped the histological phases of samples across patients. Similar to PCA analysis of epithelial pseudobulk transcriptomes (Fig. 2A, middle left panel), all *mNE* samples clustered together, separated from the majority of *mNS* samples (dendrogram in Fig. S4A); in between are clusters of the *mNS*, *M*, *D* and many *T* samples (right portion of dendrogram in Fig. S4A). Importantly, the ratio of epithelium and stromal cell abundance was a major determinant of the large clusters (simplified heat map underneath the dendrogram in Suppl. Fig. S4A). This was further supported by the observation that variation in gene expression was primarily driven by changes in composition of the various epithelial cell types rather than gene expression (cell phenotype) change, since the principal coordinates in Fig. 2A correlated with cell type proportions across samples (Fig. S4B).

Analysis of specific cell type proportions at each disease phase provided additional insights (Fig. S4C). Thus, *mNS* and *M* samples tended to contain fewer immune cells. Conversely, the *T* samples exhibited a significantly higher proportion of macrophages (1.9-fold) and T cells (1.5-fold) compared to mNE (Fig. S4 C-D). Interestingly, this feature further set *T* apart from *M*, a difference not exposed by bulk gene expression profiling of immune cells (Fig. 2A). Thus, cell type abundance, used as a feature for clustering, could extract gene→ *T*. As expected, we observed a decrease of tissue-specific transcripts with increasing malignancy, in line with ric patterns despite sample-specific variability.

### Changes in the individual cell types and their subtypes during progression

We first examined the epithelial cells and changes of the transcriptional states with progression (*mNE*, *mNS*) → *M* → *D* either differentiation arrest or dedifferentiation in neoplasia which are now considered hallmarks of malignancy^31^. The loss of differentiated cells was specifically manifest in the reduction of expression of differentiation markers, e.g., MUC5AC, FCGBP, CLCA1, MUC6 and KRT20 at the transition from M to D/T (Suppl. Fig. S4E). Thus, gastric and intestinal differentiation expression programs "faded" in the development from *M* to *D* and *T* as previously observed^7^ .

Among tumor microenvironment cells, the relative proportions of the 6 coarse-grained cell type categories and corresponding transcript profiles changed during progression (Fig. 3). Within each of these 6 groups (fibroblasts, myofibroblasts, endothelial cells, T/NK cells, myeloid cells and B/plasma cells), we also analyzed the finer-grained cell subtypes (e.g., venous, arterial, capillary and lymphatic endothelial cells) by sub-clustering these groups individually into subgroups and quantifying changes of their abundance across disease progression phases (Methods).

**Figure 3.**
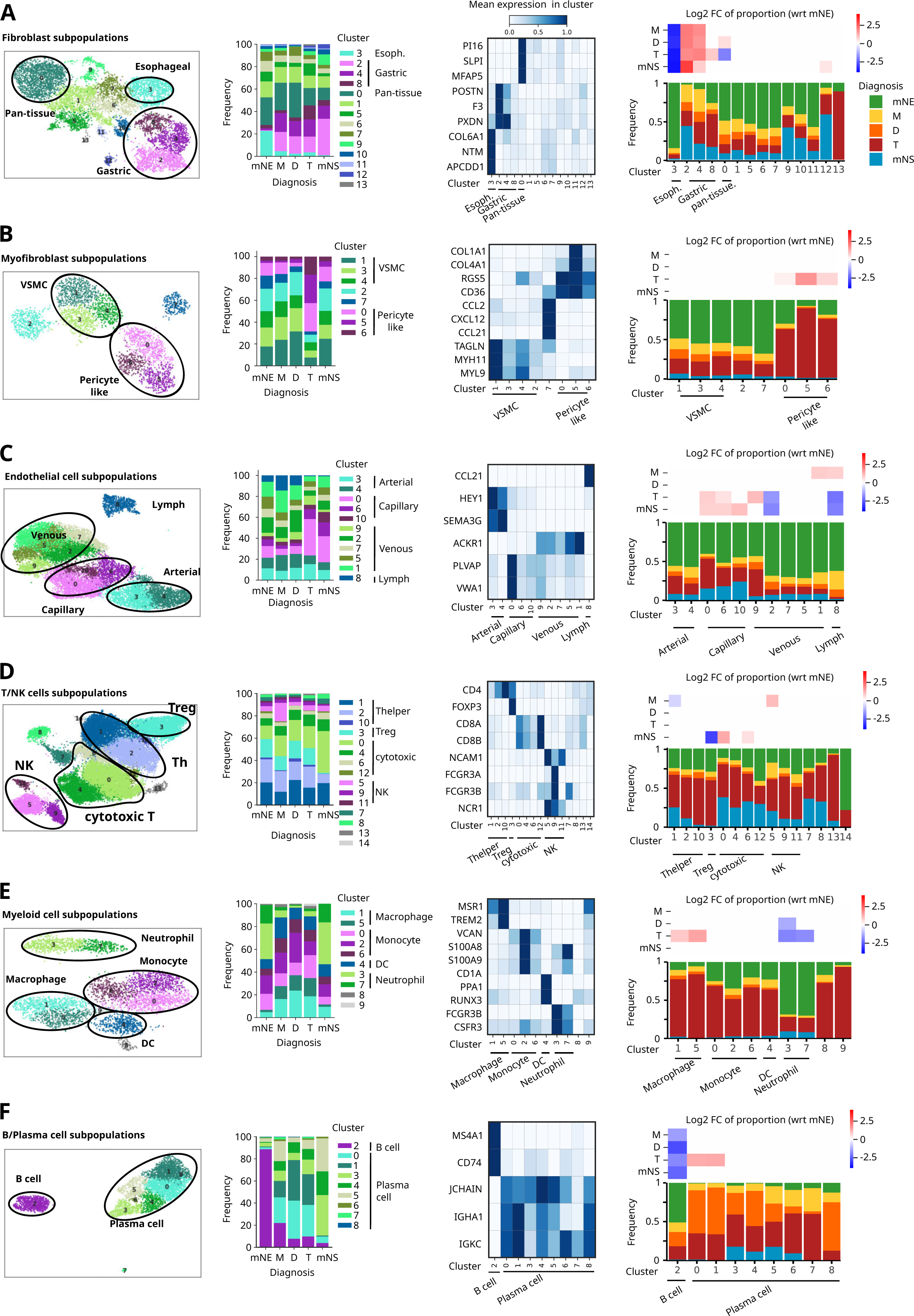
Stromal and immune cell subtype composition changes during BE progression. **(A)** fibroblasts, **(B)** myofibroblasts, **(C)** endothelial cells, **(D)** T/Natural Killer (NK) cells, **(E)** myeloid cells (macrophages, monocytes, dendritic cells (DCs) and neutrophils), and **(F)** B/Plasma cells. **First column panels:** UMAP of cells from the respective cell type (including all diagnoses), color-coded by transcriptional cluster. **Second column panels:** Composition of cells with respect to cluster (subtype) membership, for each “diagnosis” (=phase of progression)/tissue with same color codes as in UMAPs. **Third column panels**: Selected marker genes for clusters of interest. **Fourth Column panels**: TOP subpanel: Statistically significant enrichment (red) or depletion (blue) of a cellular subtype at each diagnosis (compared to *mNE*, determined by scCODA, Methods). BOTTOM panel: Composition of cells with respect to the diagnoses, for each transcriptional cluster (subtype).

**Figure 4.**
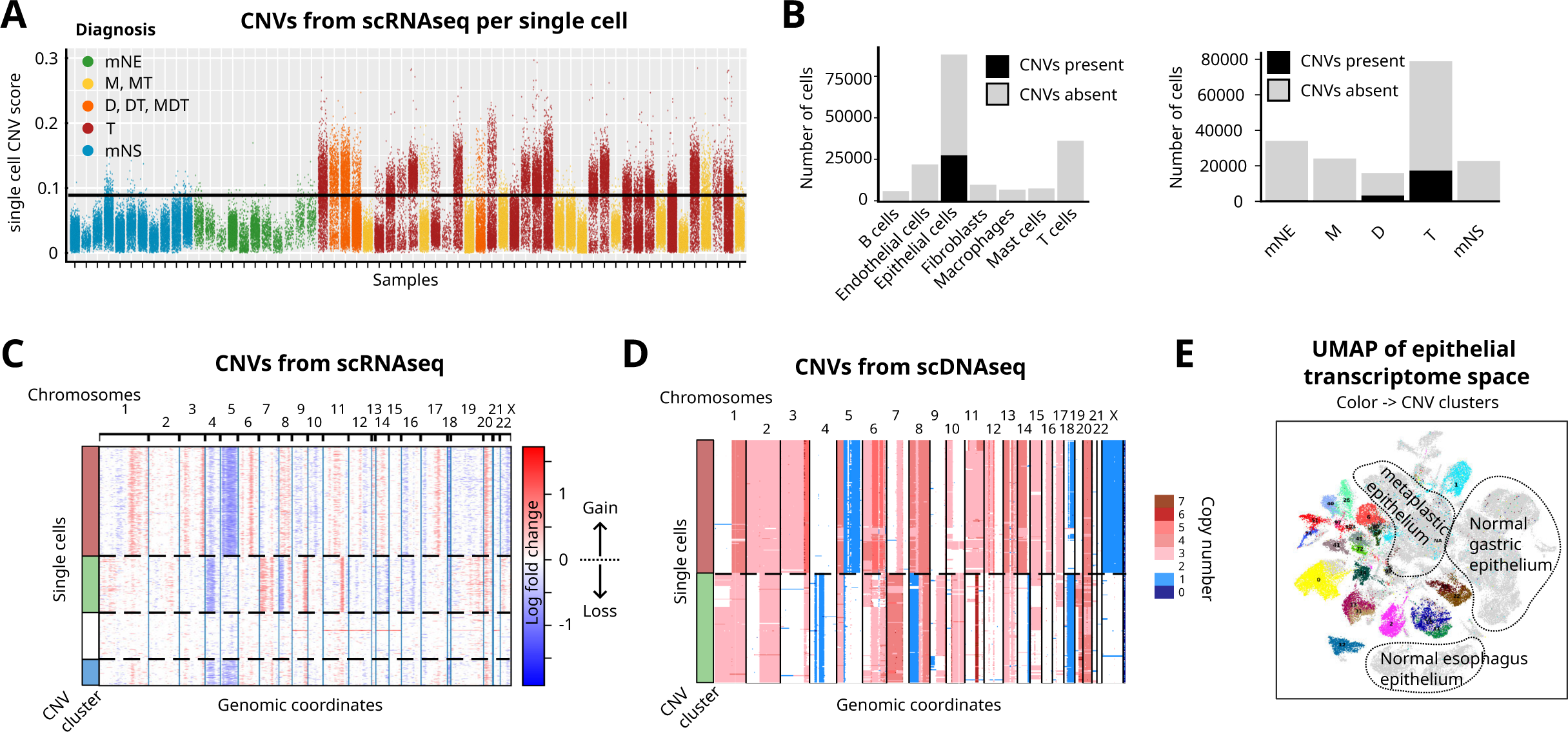
Large CNVs are markers of BE progression to malignancy. **(A)** For each epithelial cell of each sample (horizontal axis) representing the phases of progression (color labels), the CNV burden (vertical axis) was estimated based on inferred CNVs from scRNAseq (Methods). Most tumor samples contained cells with significantly increased CNV score compared to the *mNE*/*mNS* samples (black horizontal line: 99% quantile of CNV score in mNE/mNS cells). **(B)** Number of cells with (black) and without (grey) CNVs, according to cell type (left panel) and to disease progression phase (right panel). Large CNVs were restricted to the epithelial compartment and appeared at the dysplastic phase. **(C,D)** CNVs for tumor E21 inferred from scRNAseq (C) and were consistent with CNVs obtained from scDNAseq (D). **(E)** UMAP of epithelial cells in transcriptome space with each cell’s CNV cluster membership identified in (C) shown in color (grey: CNVs absent). CNV clusters closely mimic transcriptional clusters.

First, within the fibroblasts (marked by DCN and PDGFRA expression), significant changes in cell type proportions were observed (Fig. 3A). Further sub-clustering fibroblasts revealed a subpopulation differentially expressing APDDC1, NTM and COL6A1 that was highly abundant in *mNE* but almost absent in other tissue or progression phases (cyan cluster 3, Fig. 3A). Thus, an esophagus-specific fibroblast population was already lost along with the loss of squamous epithelium at the *M* stage. Correspondingly, the replacement of squamous epithelium by columnar epithelium in BE was accompanied by replacement of APDDC1, NTM and COL6A1-expressing esophagus-specific fibroblasts by PXDN, F3, POSTN-expressing fibroblast subpopulations (clusters 2, 4 and 8, Fig. 3A). These fibroblast subpopulations were absent in *mNE* but appeared in *M*, *D* and *T* tissues. Intriguingly, these fibroblasts that were associated with progression beyond *mNE* were transcriptionally almost identical to those found in matched normal stomach (*mNS*) samples. These fibroblasts would be considered to be CAFs (absent in mNE, present in T). Yet they were not “abnormal” *per se* but rather represented a cellular reprogramming to a state very similar to that utilized by the non-neoplastic nearby gastric tissues (*mNS*). We also confirmed a previously reported subpopulation of PI16-expressing “pan-tissue” fibroblasts (cluster 0, Fig. 3A)^32^ that appeared to be stable across all specimens and progression phases.

Next, we examined the myofibroblast compartment (expressing ACTA2, PDGFA and NOTCH3). We found subpopulations resembling vascular smooth muscle cells (clusters 1, 3 and 4, Fig. 3B) ^33^, pericyte-like cells characterized by COL1A1, COL4A1, RGS5 and CD36 expression (clusters 0, 5 and 6, Fig. 3B) and a cell subpopulation expressing pro-inflammatory cytokines, such as CCL2, CCL19, CCL21 and CXCL12 (cluster 7, Fig. 3B). Proportions of these subpopulations remained stable between *mNE*, *mNS*, *M* and *D* samples. The number of pericyte-like cells increased substantially in *T* samples (3.5-fold compared to *mNE*), a trend also observed in in gastric cancers^34^. This stromal subtype however was not tumor-specific, as it was also found in all other stages albeit in relatively small amounts. While neovascularization in tumor tissues produces abnormal vasculature devoid of pericytes, signaling from these mural cells to endothelial cells is thought to be critical for promoting tumor angiogenesis ^35, 36^ and, after injury, pericytes have been shown to detach from endothelial cells and transition to pro-tumorigenic, inflammatory myofibroblast-like cells^37^ .

Within the endothelial compartment, we identified four prominent subpopulations, corresponding to arterial (compared to other endothelial subclusters, differentially expressing HEY1 and SEMA3G), venous (ACKR1), capillary (VWA1, PLVAP) and lymphatic (CCL21) endothelial cells^38–40^ (Fig. 3C). Whereas all subpopulations were present at each progression phase, we observed a significantly higher proportion of capillary endothelial cells (EC) in *T* and *mNS* compared to *mNE*, again revealing a change in tissue identity during EAC progression that shifts the entire tissue, as a unit, towards the phenotype of matched gastric tissue (*mNS*) (Fig. 3C). Furthermore, the frequency of a particular subpopulation of venous endothelial cells (VEC, cluster 2, Fig. 3C) and lymphatic endothelial cells were also lower in *T* and in adjacent *mNS*. VEC are the cellular source for capillary endothelium during angiogenesis^41^, and their reduction along with the increased number of capillary EC may reflect depletion of VEC due to ongoing angiogenesis in the tumor. Similarly, the low number of lymphatic endothelial cells in *T* is in line with the general absence of lymphatics within tumors (although tumor-marginal lymphatics are critical for lymphatic metastasis)^42, 43^ . However, since these vasculature remodeling features seen in *T* were also found in the non-cancerous *mNS*, we cannot rule out that this may reflect an intrinsic property of gastric mucosa that distinguishes it from the squamous epithelium of matched normal esophagus that would be co-opted during EAC progression.

Marked changes of cell type composition in the immune compartment of the tumor microenvironment were revealed by single-cell transcriptomics of the various phases of progression although immune-cell specific bulk transcriptomes were not a discriminatory feature (Fig. 2A, right panel). Among T cells and NK cells, we identified cytotoxic T cells (CD8A), T-helper cells (CD4), T-regulatory cells (FOXP3) and different subtypes of NK cells (GNLY) (Fig. 3D). Most of these subpopulations were stable across the progression phases with the exception that *mNS* tissues had reduced levels of T-regulatory cells (cluster 3, Fig. 3D) and displayed slightly higher numbers of cytotoxic T cells compared to the squamous *mNE*. Interestingly, *M* samples exhibited a sizable increase in NK cells, notably of the NCAM1(CD56^high^) immunosuppressive subtype^44, 45^ (cluster 5, Fig. 3D; FCGR3A, FCGR3B, NCAM1/CD56, NCR1) and a decrease in T-helper cells (cluster 1, Fig. 3D), which jointly indicated acquisition of an immunosuppressive environment in metaplasia.

In the myeloid compartment, scRNAseq readily identified macrophages (MSR1, TREM2), monocytes (VCAN, S100A8), dendritic cells (PPA1, RUNX3) and neutrophils (FCGR3B, CSF3R) (Fig. 3E). Neutrophils were abundant in both *mNE* and *mNS* samples (Fig. 3E), but their frequency was significantly lower in *D* and *T* samples. Furthermore, as mentioned, there was a significant increase (3.2-fold) in macrophages in *T* samples (Fig. 3E). The frequencies of monocytes and dendritic cells were constant across disease progression phases.

Both B (MS4A1) and Plasma (JCHAIN, IGHA1) cells were present at all disease progression phases (Fig. 3F). Notably, *mNE* samples showed abundant B cells but contained few plasma cells. With the onset of metaplasia, the fraction of B cells decreased significantly, while the fraction of plasma cells increased substantially, a finding that has been previously associated with outcome of EAC^46^ .

### scRNA-seq analysis reveals large CNVs in the dysplastic phase and patient-unique clonal history of malignant cells

Genomic instability, manifested as copy number variants (CNVs), is a hallmark of EAC. Appearance of CNVs in BE has been considered an early biomarker for progression^10, 47^ . To both identify aneuploid cells and estimate clonality, scRNA-seq reads were used to estimate CNVs in each cell^48^ .

For all scRNAseq samples, we inferred CNVs in epithelial cells using inferCNV (Methods) with matched normal samples (*mNE*/*mNS*) as reference. As expected, tumor samples contained a substantial number of epithelial cells with CNVs (Fig. 4A-B) not found in stromal cells (Fig. 4B). Dysplastic samples contained cells with CNV profiles similar to cells from the corresponding tumor, suggesting a clonal relationship between dysplasia and tumor (Fig. S5A-B). Most tumors contained one major copy number profile, indicating that the cells were clonally related. No large-scale CNVs were seen in metaplastic samples. Only two samples diagnosed as metaplasia exhibited small amounts CNV. However, these exceptions were probably due to contamination with dysplastic and tumor cells (data not shown). Of note, any small-scale CNVs in metaplasia could be undetectable given the genomic resolution limit (>10MBp) of the methodology.

To confirm scRNAseq-inferred CNVs, we performed single-cell whole exome DNA sequencing on selected samples. Comparison of CNV profiles from E21 tumor, either inferred from scRNAseq (Fig. 4C) or obtained from the matched scDNAseq sample (Fig. 4D), revealed good agreement. Both methods detected the same major clones: the first clone was characterized by chr1.q and Chr8 copy number gain, and by chr5 deletion (red CNV cluster, Fig. 4C-D); the second clone showed copy number gains of chr7 and chr8.q and deletions of chr4 and chr8.p (green CNV cluster, Fig. 2C-D). scDNAseq did not identify a third clone (blue CNV cluster in Fig. 2C), possibly due to the limited number of cells used for scDNAseq.

The copy number profiles of each patient’s tumor were unique and manifested as distinct transcriptional clusters of epithelial cells (color-coded clusters in Fig. 4E) which dominated the interpatient heterogeneity observed in the tumor epithelial cells. In summary, our observations showed that in our small cohort, large-scale copy number variation arose at the dysplastic phase, consistent with previous findings^10, 49, 50^.

### Changes in extracellular matrix (ECM) composition during progression mirror concerted epithelial and stromal cell changes

The extracellular matrix (ECM) is central to the molecular and mechanical interaction between epithelial and stromal cells. We used mass spectrometry to examine ECM composition in a subset of the 12 patients for whom we had single-cell transcriptomic data (Fig. 1A, see Methods), focusing on comparison between mNE and T: 6 tumor samples (T) and 4 patient-matched normal esophagus samples (mNE) (Fig. S1A). In the ECM-enriched material purified from these samples, 1994 protein groups with at least two unique peptides were quantified by mass spectrometry, roughly half of which were categorized as “extracellular” proteins. Among them 78 were core matrisome proteins and 73 were matrisome-associated proteins (Fig. 5A). Principal component analysis of the samples based on the proteomic profiles showed separation between *T* and *mNE* samples (Fig. 5B).

**Figure 5.**
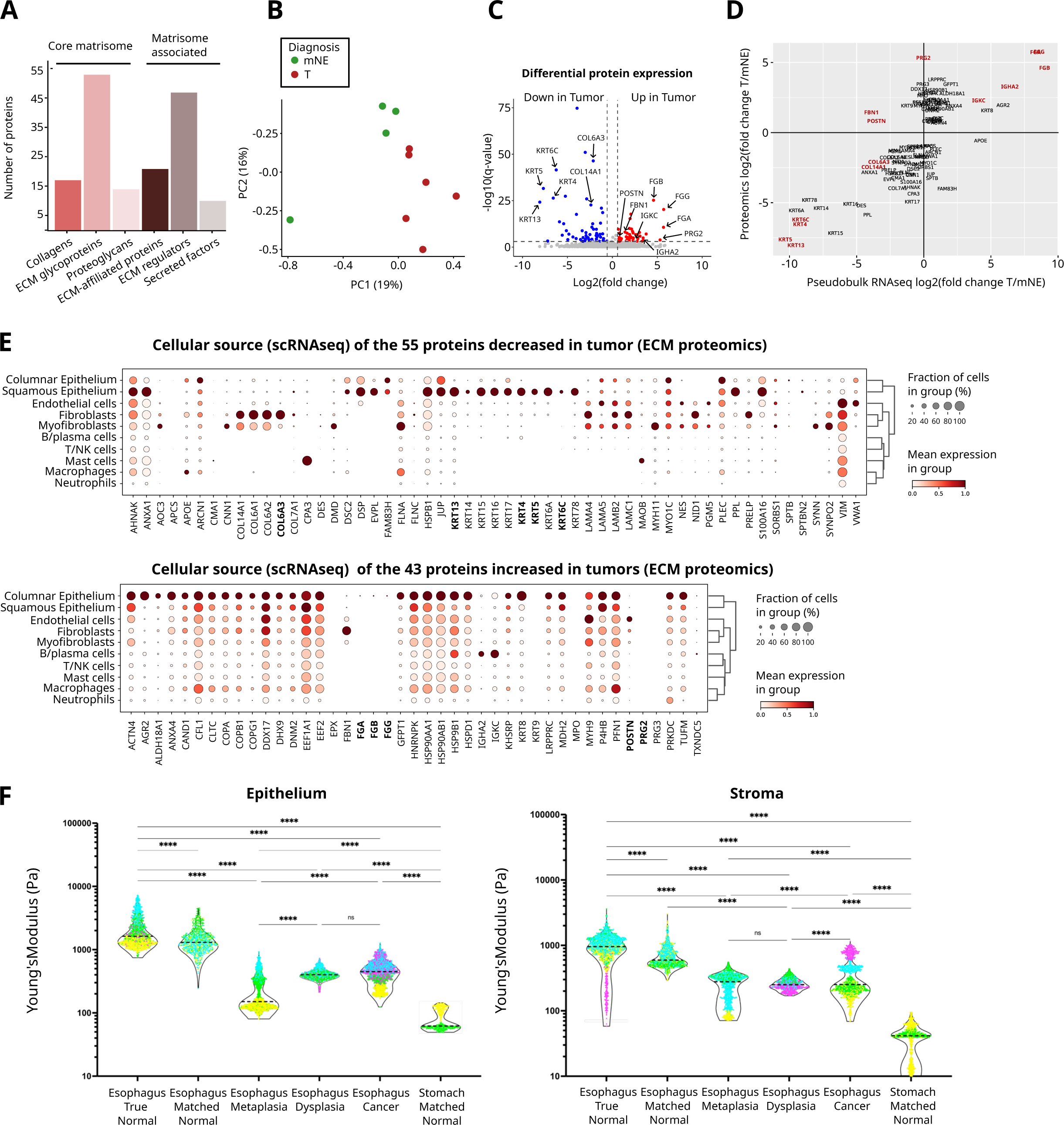
Proteomic analysis and Atomic Force Microscopy (AFM) reveal changes in the extracellular matrix composition and tissue mechanics during BE progression. **(A)** Number of matrisome or matrisome-associated proteins detected by ECM proteomics grouped by protein class. **(B)** PCA of proteomic samples (*mNE*: green; *T*: red) shows differences in ECM-enriched protein extract composition between *mNE* and *T* samples. **(C)** Differential ECM protein abundances between *mNE* and *T* samples shows 43 and 55 proteins upregulated and downregulated in *T* compared to *mNE*, respectively. Proteins with the highest up/down-regulation in *T* vs. *mNE* are identified. **(D)** Positive correlation between ECM proteomic-(*y*-axis) and respective scRNAseq (*x*-axis) fold-changes in matched samples. Proteins of interest are highlighted in red font. **(E)** Expression of significantly down-regulated (top panel) and up-regulated (bottom panel) proteins identified by proteomic analysis (see C) grouped by scRNAseq cell type (including cells from all diagnoses). **(F)** Quantification of epithelium and stroma stiffness using AFM in a distinct cohort for the phases of progression, shown as violin plots. Each color represents an esophageal sample at a different phase from a given patient; included are here also samples from tumor-free individuals (“true normal”) as opposed to the matched-normal” samples from EA patients. Donors: *n*=3, for matched normal esophagus, esophagus metaplasia, and dysplasia; n=4, for esophageal cancer and true normal esophagus; n=2 for matched normal stomach. Two-way Anova test with Tukey post hoc test was used as statistical test. Thick and thin dashed lines represent median and quartiles in the distribution, respectively. *n* = 50-150 regions per patient, ns: not significant, **** p <0.001.

Comparing pooled *T* versus *mNE* samples, we found 98 proteins with significantly altered abundances in the ECM-enriched extract (q-value < 0.001 and |fold change| > 1.5). 55 of them showed decreased and 43 increased expression in *T* compared to *m*NE (Fig. 5C). The 10 proteins with the most dramatic changes (9 “down-regulated” and 1 “up-regulated”), as well as a 22-protein matrisomal core protein signature, are presented in Fig. S6. The decrease in ECM protein expression during progression from *mNE* to *T* was predominantly represented by some specific collagens, in line with the reduction in collagen-producing fibroblasts in *M*, *D* and *T* compared to *mNE* observed in the scRNAseq dataset (Fig. 3A, fibroblast cluster 3)^51^. As expected, expression of keratins associated with the squamous *mNE*, also decreased during progression to *T*. Conversely, *tumor associated* ECM had increased levels of proteoglycans PRG2 and PRG3 as well as fibrinogens FGA, FGB and FGG (Table S6).

### Integration of scRNAseq and proteomics reveal cellular sources of ECM protein changes

We next examined whether scRNAseq findings were consistent with those of ECM proteomic analysis for the 10 patient samples analyzed by both methods (Fig. 5D). scRNAseq could provide information about the cellular source(s) of the 98 ECM-associated proteins (Fig. 5E) with significantly altered abundance in *T* vs *mNE*. As Fig. 5D shows, relative ECM protein abundances (fold-change in *T* vs. *mNE*) for this set of proteins (Fig. 5C) correlated well with the fold changes of transcripts computed as pseudo-bulk expression from the equivalent patient samples (Spearman R=0.73).

Specifically, scRNAseq linked changes in protein abundances at the tissue level to shifts in specific cell types/subtypes that produced them. For instance, mRNA expression of collagens 6A1, 6A2 and 6A3 was reduced in *T* samples, a phenotype that was consistent with the loss of fibroblast subtypes that strongly expressed transcripts encoding these same ECM molecules during progression (cluster 3, Fig. 3A). The marked reduction in collagen 14A1 in the ECM proteomic analysis could also be explained by the observed shift in fibroblast subtype fractions, as transcripts for COL14A1, while expressed in *mNE* fibroblasts, were no longer expressed in matched normal stomach (mNS) fibroblasts, the fibroblast subtype that became predominant at the *M* phase and subsequent *D* and *T* phase (Fibroblast clusters 2, 4 and 8, Fig. 3A).

Notable departures from the correlation between ECM proteomics and scRNAseq included fibrillin (FBN1) whose protein abundance increased but whose transcript levels decreased instead. This discordance might be explained by its predominant expression in adipose tissue, a tissue compartment represented in the proteomic analysis but totally absent in single-cell transcriptomics because adipocytes are lost during the cell dissociation process^52^. Similarly, the relatively high abundance of the two proteoglycans, PRG2 and PRG3, in tumor ECM was not concordant with transcript analysis which showed no expression in *T* and *mNE*. This finding is consistent with the predominant production of these proteoglycans in liver and bone marrow (and in female reproductive organs) and suggest that these proteins may have been deposited via circulation in the esophagus tumor stroma^53^ . Similarly, fibrinogens FGA and FGB, which are solely produced in the liver, were also unexpectedly elevated in the tumor ECM proteome of several patients but not in the corresponding tumor transcriptome ^54^, pointing again to the synthesis of fibrinogen in the liver and subsequent deposition at the tumor site. Strikingly, concordance between fibrinogen proteome and transcriptome was observed for one patient (E14). This concordance between fibrinogen protein and local transcripts could be explained by the rare but previously reported occurrence of hepatoid differentiation (documented by identification of an alfa-fetoprotein (AFP)-expressing cluster) of the EAC of this patient.

Examining cellular sources confirmed that one major protein absent in the tumor ECM-enriched extract, KRT13 (Fig. 5C, Suppl. Table S6), a keratin filament protein characteristic of stratified squamous epithelium, resulted from the replacement of the esophagus squamous epithelium by a gastro-intestinal columnar epithelium (Fig. 5E). Another protein of interest was periostin (POSTN), an ECM protein that was moderately but significantly increased in the tumor ECM (Fig. 5C, Table S6). Cell-type specific transcript analysis (Fig. 3A) had shown that it was a prominent marker of a subtype of fibroblasts associated with gastric tissue, and characteristic of progression from mNE to *M*, *D*, and *T* (fibroblast clusters 2, 4 and 8, Fig. 3A). Tumor transcript analysis in the pseudo-bulk data (Fig. 5D) failed to detect an increase in transcripts of POSTN in this comparison, although it has been previously implicated in BE^15, 55^ . However, single cell-resolution analysis showed that, while POSTN transcripts were elevated by almost two-fold in tumor-associated fibroblasts, in this same sample set, the tumor capillary endothelial cells (Fig. 3C), showed instead a down-regulation of POSTN by more than two-fold (Fig. S7A-B). Thus, a shift of transcript levels for a given gene in opposite directions in distinct cell types/subpopulations might conceal differential expression in bulk RNAseq (Fig. S7A-B).

Finally, among the most significantly increased non-ECM proteins in the ECM-enriched preparation of T compared to mNE samples (Table S6), were Ig-related subunit chains IGHA2 and IGKC (Fig. 5C). This increase could readily be attributed to the surge of plasma cells in *T* compared to *mNE*, consistent with the cell-type composition changes noted above (Fig. 3E)^46, 56, 57^.

Taken together, scRNAseq and cell type-level resolution analysis not only allowed us to trace back the cellular origin of changes observed in bulk analysis methods, such as proteomics, but also allowed us to expose opposing expression changes in distinct cell type compartments that might have masked a cell-type specific differential expression.

### Changes in mechanical stiffness during progression confirm concerted epithelial and stromal alterations

Since ECM changes have been linked to altered mechanical tissue stiffness, which in turn can affect tumor-promoting signaling^58^, we used atomic force microscopy (AFM) to assess the stiffness of esophageal tissues during disease progression by comparing AFM characteristics of samples across disease progression phases for epithelial and stromal stiffness (Fig. S8A-B). Stiffness measurements of epithelium were found to be in a range similar to those previously published (0.6-3.0 kPa)^58–60^ (Fig. 5F). Strikingly, stiffness of the *mNE* (squamous) epithelium was ∼20-times higher than that of *mNS* (columnar) epithelium (1.2 vs. 0.06 kPa).

Metaplastic Barrett’s esophagus (*M*) not only acquired a gastric histology but also a “softer” gastric-like stiffness. However, as expected, this lower gastric-like stiffness increased with pathological progression. Although the stiffness of epithelia in the columnar lesions *M*, *D*, *T* was 3-8 times lower (0.15, 0.4 and 0.45 kPa, respectively) compared to that of the squamous *mNE*, stiffness of these gastric-like columnar lesions was still 3-8 times higher than that of normal tissue, *mNS* (Fig. 5F). Thus, while the esophageal columnar lesions adopted a softer gastric-like identity, their stiffness was still significantly higher than that of the non-cancerous *mNS.* This observation is in accordance with the well-documented general concept of increased stiffness in diseased/cancer tissue compared to healthy tissue^61^.

AFM analysis also revealed mechanical changes in the stroma. Consistent with the difference in epithelial type, stiffness of the stroma of *mNE* esophagus (∼0.6 kPa) was dramatically higher than that of *mNS* (∼0.04 kPa) (Fig. 5F). However, in most samples, the stroma of the columnar lesions *M*, *D*, *T* was similar or only slightly softer than that of *mNE* (0.24-0.26 kPa vs. 0.6 kPa), with large variation between samples; some regions exhibited much higher stiffness, reflecting the general trend of neoplastic tissues (Fig. 5F). The moderate decrease in stromal stiffness that correlates with replacement of squamous by columnar tissue during progression was in accordance with the observed decrease in ECM proteins, such as laminin and prolargin, as well as stromal cell cytoskeletal proteins that link cell mechanics to the ECM (e.g. desmin, calponin, dystrophin, and filamins A and C), as identified by the ECM proteomics (Table S6).

### Multiplexed protein imaging reveals reorganization multi-cellular tissue neighborhood during progression

To interrogate the spatial relationships of the cell types within the progression phase of EAC, we performed CODEX multiplexed protein imaging on 27 imaging regions in the esophageal tissue from 5 of the 12 patients (Fig. 6A, Fig. S9A)^62–64^ . Diagnosis of tissue and phase of progression (*mNE*, *M*, *D*, *T*; *mNS*) was verified by pathologists with regard to proportions of cells using H&E (hematoxylin and eosin) staining of the same sections post-CODEX (Fig. S9B-D, Table S8). Given our focus on stroma, we designed our 54-antibody panel to interrogate epithelial, stromal and immune cell types (Table S9). The CODEX marker panel allowed us to identify 45 unique cell types (Fig. S9E), 25 of which were epithelial ^7^, including “squamous” (*mNE*), “foveolar” (*M*), p53+ (p53-expressing) and lineage-negative epithelial cells (*D*, *T*) (Fig. 6B,E, Fig. S9F), which were consistent with the overall change in cell type proportions seen in the scRNAseq (Fig. S10).

**Figure 6.**
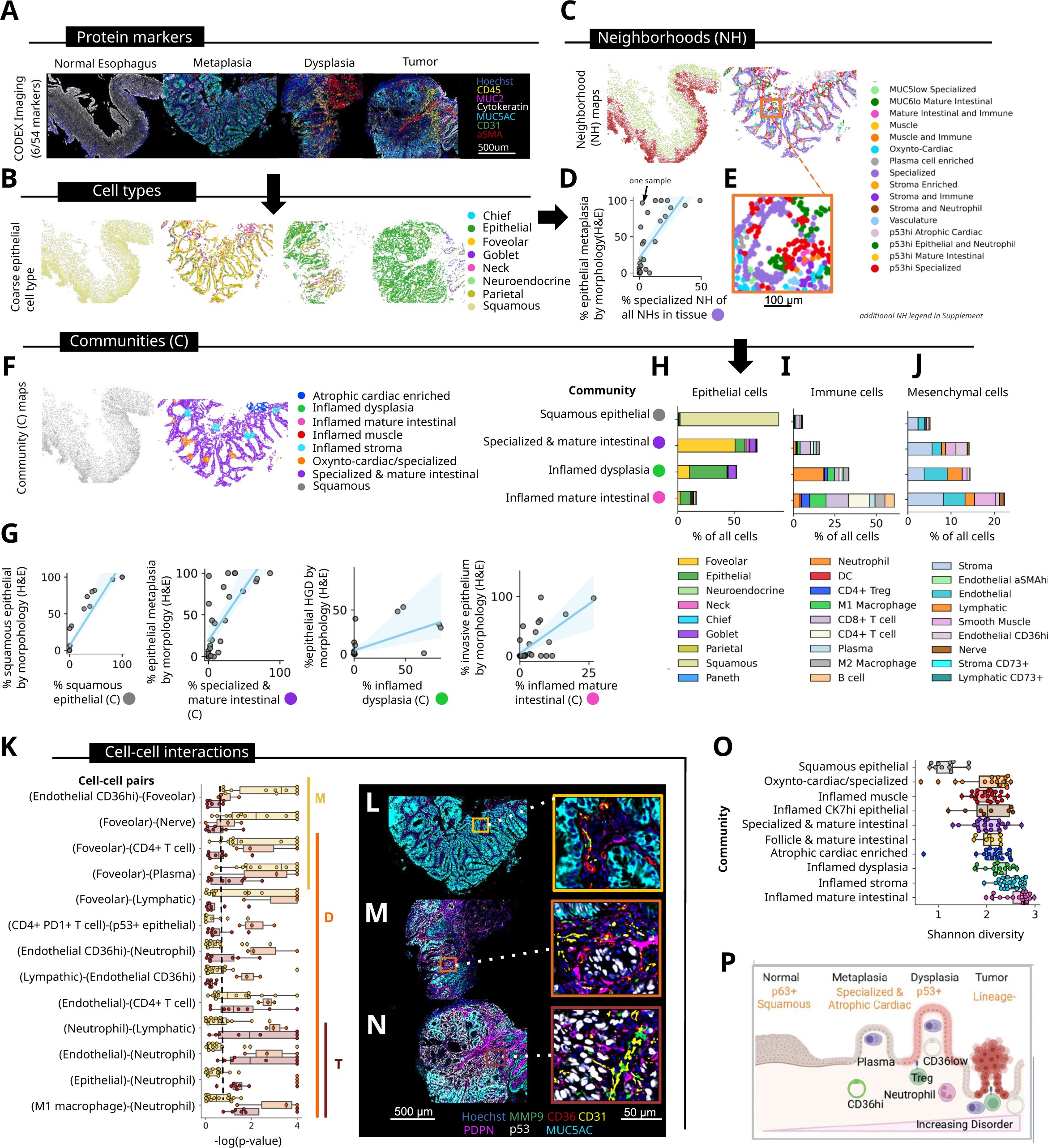
CODEX multiplexed proteomic imaging identifies epithelial and stromal tissue structures, exposing cell state changes. **(A)** Representative images of *mNE*, *M*, *D* and *T* esophageal tissues from one patient (of a total of 5) from CODEX multiplexed fluorescent imaging for 6 of the 54 markers queried (scale bar = 500 um) and corresponding **(B)** coarse-grained, CODEX-defined epithelial cell types mapped back to tissue coordinates. **(C)** Neighborhood (NH) maps for *mNE* and *M* . **(D)** Percentage of metaplasia, as determined by a pathologist (*y*-axis) versus percentage of the “Specialized” NH from CODEX analysis of each of the 27 regions imaged (*x*-axis). **(E)** Magnified region of NH map for *M* sample shown in C (scale bar = 100 um). **(F)** Community (neighborhood of neighborhoods) maps for representative *mNE* (left panel) and *M* (right panel) samples. **(G)** Percentage of squamous (*mNE*) esophagus, metaplasia (*M*), high grade dysplasia (HGD), and invasive EAC epithelium as determined by a pathologist (*y*-axis) versus percentage of community determined by CODEX for each of the 27 regions imaged (*x*-axis). **(H-J)** Cell type composition for each of the 4 communities (*mNE*: grey; *M*: purple; *D*: green; *T*: pink) that correlated with diseased epithelial in panel G, broken down by (H) epithelial, (I) immune, and (J) mesenchymal groupings. Percentages are provided in Suppl.Table S10. **(K)** cell-cell interaction analysis with -log(p-value) for each cell type pair compared to 10,000 random permutations, colored by overall disease state. **(L-N)** Representative images of the phases or progression: *mNE* (panel L), *D* (M), and *T* (N) from one donor (5 donors total) from CODEX imaging for 6 of the 54 markers queried (scale bar = 500 um) and of magnified regions (scale bar = 50 um). **(O)** Shannon’s Diversity for cell types and proportions within each CODEX-defined community. **(P)** Schematic representation of overall changes in epithelial and stromal connections.

Because CODEX preserves tissue geography, we next asked if spatial proximity of cell types to one another was altered during disease progression in light of the dramatic changes in cell type composition observed by scRNAseq (Figs. 2 and 3). Thus, we performed cellular neighborhood (NH) analysis to identify conserved multicellular microstructures (Methods)^65^ and expose changes in tissue organization not observable by shifts in global cell proportions using scRNAseq. We identified 24 unique multicellular neighborhoods that were labeled based on enrichment of cell types within each neighborhood (Figs. 6C and S11A-B). For example, as expected, *mNE* samples were highly enriched for an “*apical squamous”* NH - characterized by Annexin A1+ squamous cells - and for a “*basal squamous”* NH - characterized by both Annexin A1+ and p63+ squamous cells (Fig. S11C,D).

We also detected a NH enriched for foveolar and goblet cells (Fig. S11B, F-G) that we termed "*specialized”* NH, consistent with previously defined gland phenotypes in metaplasia^66^ . As expected, this *“specialized”* NH correlated with the percentage of metaplastic epithelium (Fig. 6D, Fig. S12A). We also detected NHs that were consistent with known epithelial organization (e.g., *“mature intestinal”* NH, “*oxynto-cardiac”* NH, “*atrophic cardiac”* NH). In addition, we identified novel conserved NHs (e.g., “*muc5low specialized”* NH*, “muc6low mature intestinal”* NH) characteristic of BE epithelial organizations (Figs. S12A and S13A). Analysis of neighborhoods allowed us to assign specific molecular alterations of malignant cells to this higher-level pattern of tissue organization. For instance, the “*p53hi atrophic cardiac”* NH correlated with the proportion of metaplastic epithelium, attributing p53 expression to particular glands (Figs. S12A and S13A).

In the stroma, we identified several consistent neighborhood organizations (e.g., a “*stroma and immune”* NH*, a “stroma and neutrophil”* NH*, and a “follicle, smooth muscle, vasculature”* NH). Interestingly, presence of the “*stroma and neutrophil”* NH correlated with the proportion of epithelium classified independently as high-grade dysplasia by two expert pathologists across all samples (Fig. S11E). This NH also was enriched for CD4+ T cells, lymphatic and vascular endothelial cells, and antigen-presenting cells (Fig. S11B).

### Communities of cell neighborhoods illustrate rearrangements of epithelial-stromal cellular entities during progression

Multiple NHs correlating with the specific histological phase of BE progression (*M*, *D*) suggested that the size of an effective cellular neighborhood structure was smaller than the larger overall pathology that underlies the histological diagnosis. This feature is reflected in the unique NH calls for cells of a single gland and in the local stromal NHs for a given histological pathology (Fig. 6E). To better align multicellular structures to the overall histological diagnosis, we thus evaluated the co-localization of multiple NHs, referred to as “communities” (of neighborhoods). We defined specific communities of NHs by increasing the number of neighborhood-defining cells (100), using the NH labels as input^67^ . This process revealed a total of 10 unique communities (C) of neighborhoods (Fig. S12B-C). Thus, the *“apical squamous”* NH and *“basal squamous”* NH were found within a single “*squamous epithelial community (C)*” because of their characteristic occurrence in proximity to each other (Figs. 6F & S12B). The presence of the “*squamous epithelial C”* correlated across all samples with the percentage of squamous epithelium (Fig. 6G). Three other communities of NH showed positive correlations with the proportion of epithelium: *“Specialized and Mature Intestinal C”* with M, *“Inflamed Dysplasia C”* with D and “*inflamed mature intestinal C”* with T (Fig. 6G).

The relationship of community structures with the phases of progression afforded a new looking glass for considering tissue geography in the cellular changes associated with BE progression by analyzing cellular composition of communities instead of the global composition as captured by scRNAseq (Fig. 6H-J, Table S10). Overall, this analysis revealed an increase in immune and mesenchymal cells in the communities with progression.

Within the epithelial cell compartment, with progression the squamous cell community (*"squamous epithelial C”*) yielded to the foveolar community (*“Specialized and Mature Intestinal C”*) that increased in epithelial cells that were negative for squamous and metaplastic lineage markers (Figs. 6H, S9E & S12D) reflecting changes both at the level of glandular structures, towards gastro-intestinal formations, as well as loss of cellular differentiation associated with progression.

Within the immune cell compartment, the proportion of neutrophils increased within the “ *inflamed dysplasia C”* (Figs. 6I & S12E). Indeed, three of the neutrophil-enriched NHs were specifically enriched within the “*inflamed dysplasia C”* (Fig. S12F). Strikingly, while most immune cell types also increased within the “*inflamed mature intestinal C”*, which correlated with invasive epithelium or tumor pathology, neutrophils did not. Instead, macrophage subsets were enriched (Figs. 6I & S12G). This “*inflamed mature intestinal C”* was enriched with diverse NHs, namely the “*mature intestinal and immune”* NH, the “*APC enriched immune”* NH, and the “*stroma and immune NH”* (Fig. S11B). Each of these NHs were characterized by abundance of immune cells (e.g., CD4+ T cells, macrophages, DCs, CD8+ T cells) and absence of neutrophils. Finally, in this *“inflamed mature intestinal C”*, the proportion of CD4+ Tregs also increased (Figs. 6I, S12I & S13H). This trend may reflect the role of Tregs in preventing neutrophil accumulation during tissue repair^68^.

Analysis of mesenchymal cells in communities revealed that the CD36hi endothelial cell population was robust in the “*specialized & mature intestinal C” (associated with metaplasia)* but reduced in the “*inflamed dysplasia C”* and “*inflamed mature intestinal C*” (Fig. 6J). Interestingly, the abundance of CD36hi endothelial cells within all samples negatively correlated with the abundance of CD4+ Tregs (Fig. S12J). This negative correlation was in accordance with the opposite phenotypes of a CD36hi anti-tumorigenic state and a CD4+ Treg high immunosuppressive, pro-tumorigenic state. This finding underscores the added value of evaluating cell proportion changes with structural guidance of cell communities and is consistent with the downregulation of CD36 in chronically-inflamed tumor stroma^69^ .

### Cell type combinations in cell-cell interactions in stroma reprogramming during progression

We next exploited the spatial NH information to examine cell-cell interactions (Figs. 6K & S13A-C, Table S11). To evaluate the enrichment of cell-cell interactions in particular progression phases (*M*, *D*, and *T*), we calculated the frequency of neighbors using a nearest neighbor approach and compared the frequency of occurrences to null-models, achieved by 10,000 permutations of cell type locations (Methods). Cell-cell interactions shared by all phases included those between immune cells (CD8+ T cell, “M1” macrophage, and plasma cell) and stroma cells (Fig. S13D). This was consistent with the increase in plasma cells across all disease phases observed with CODEX (Fig. S13E) and scRNAseq (Fig. 3F). Plasma cells are normally seen within mucosal areas of the intestine and form a conserved niche^67^ . Thus, the increase in plasma cells (Fig. 6I) and the identification of a specific “*plasma cell enriched”* NH (Fig. S11B) may represent an aspect of reprogramming of the stromal environment.

Many unique interactions involving foveolar cell type were enriched in BE (Fig. 6K). For instance, in *M*, but not *D* or *T* samples, foveolar cells were found next to the CD36hi endothelial cells and to nerves (Figs. 6K-L & S13F), in line with their enrichment within the “*specialized & mature intestinal C*” (Fig. 6I). Additionally, in both *M* and *D*, foveolar cells were found close to CD4+ T cells and plasma cells (Figs. 6M-N & S13G-H), consistent with plasma cell niches known to accompany intestinal epithelial transitions. Finally, we observed that neutrophils paired with lymphatic cells, endothelial cells and M1 macrophages in *D* and *T* (Figs. 5M-N & S13I).

### Epithelial and stromal cellular communities become increasingly diverse during progression

Communities associated with BE progression had altered proportions of both epithelial and stromal cell types (Figs. 6H-J & S12C-G, Table S10). Indeed, the diversity of gland structures has been shown to be associated with progression in BE^70^ . However, our analysis also took into account the stromal component. We quantified the diversity of cell types in communities using the CODEX markers (Fig. 6O) and found an increase in Shannon’s diversity index *H* in the communities associated with progression: *H*(“*squamous epithelial C”)* < *H*(“*specialized & mature intestinal C”*) < *H*(“*inflamed dysplasia C”*) < *H*(“*inflamed mature intestinal C”)* (Fig. 6O). This pattern indicated that areas with high (chronic) inflammation and lacking well-differentiated structures were more likely associated with invasive tumors.

## DISCUSSION

In this study, we describe concomitant epithelial and stromal changes occurring during progression from a premaligant state (BE) to invasive cancer (EAC). This was made possible by analyzing 64 tissue samples collected from 12 individual patients that represent the sequential histological phases of BE progression from matched “normal” esophageal tissue (*mNE*), to Barrett’s metaplasia (*M*), dysplasia (*D*) and invasive adenocarcinoma (*T*) in the same patients. The matched normal stomach tissue (*mNS*) constituted a key reference because BE is the adaptive acquisition of a gastric/intestinal-like tissue. The disease progression phases co-occur within the same patient and thus can, to some extent, be regarded as a progression trajectory. Our data show marked phase-specific alterations shared by the twelve patients at multiple size scales, with respect to transcriptome, cellular content, large-scale CNVs, ECM proteomic profile, tissue mechanics, and histological architecture, with some patient-specific variability. Our studies underscore the importance and highlight new opportunities of not only multi-omics but also multi-scale integrated analysis.

A major insight is that, despite absence of obvious histological alteration in the stromal compartment, the stroma also underwent, from early in progression (*M* phase), drastic changes in cellular, mechanical and molecular identity and tissue organization that complemented those seen in the epithelium. The multi-scale approach enabled a combined assessment of change at the levels of cell numbers (cell type proportions) and cell states (molecular cell subtype) which affected epithelial cells, fibroblasts, endothelial cells, immune cells, acting in concert during advancement to malignancy and revealed wide-spread and coordinated non-genetic cell plasticity.

A key finding was the early loss of esophagus-type and gain of gastric-type fibroblast subpopulations with onset of epithelial metaplasia (BE) and in ensuing progression. Thus, the epithelial change from squamous to columnar (*mNE* → *M*) was accompanied by a commensurate transition from squamous-supporting fibroblasts to columnar-supporting fibroblasts (Fig. 3A); this shift was robust enough to separate *mNE* from *mNS*/*M*/*D*/*T* in PCA space based on pseudo-bulk transcriptomes of stromal cells (Fig. 2A). This result is consistent with concerted changes of epithelium and stroma as one unit in the tissue trajectory of healthy to premalignant and malignant tissue. Besides acquiring a gastric identity upon transition to *M* and later to *D* and *T*, these fibroblasts also acquired characteristics previously documented in carcinoma-associated fibroblast (CAF) populations, i.e., expression of TGF-beta targets ^71^ . Most investigators believe that tumor cells program fibroblasts into pro-tumorigenic CAFs. However, substantial data suggest that pro-tumorigenic fibroblasts may already be present before development of a tumor^69, 72^ . Our analysis of these esophageal samples at single-cell resolution confirmed this observation. The origin of these pro-tumorigenic fibroblasts is intensely speculated. The presence of pro-tumorigenic fibroblasts in BE may simply reflect the outgrowth of gastric tissue as one epithelium-stromal unit or it may involve the migration of gastric fibroblasts. Alternatively, presence of the latter may reflect a cell-level state transition (i.e., transdifferentiation) of local esophageal fibroblasts originally supporting the squamous epithelium during progression or a reprogramming of progenitor (fibroblast) cells. Finally, recent studies have demonstrated the transition of injured pericytes to a CAF-like state that are pro-tumorigenic^37^ .

The shift in fibroblast subpopulation occurred early at the *mNE*→*M* transition (Fig. 3A). This might entertain the interpretation that stromal changes precede and possibly drive epithelial transformation to malignancy^73^ . Ongoing analyses on patients diagnosed with BE but who, unlike the patients in his study, have not progressed to dysplasia or carcinoma will allow us to discriminate between events already occurring at the BE disease phase in the absence of malignancy from events occurring in a BE which has been subjected to a potential tissue field effect by the neighboring tumor. It should also be noted that tissue plasticity described at these early stages of disease occurs in the absence of major mutational or genomic alterations and thus represent examples of non-genetic plasticity in all tissue components.

To more explicitly demonstrate the added value of single-cell resolution compared to bulk (whole sample) analysis and explore a multi-scale analysis, we generated pseudo-bulk transcriptomes from the scRNAseq data and compared samples as wholes or selectively, with respect to specific cell types. This analysis showed that, at the whole tissue level, clustering of histology-based diagnosis (disease phases) and corresponding disease progression phases was most obviously associated with changes in the epithelium. However, extending previous reports emphasizing the role of stromal and immune cells in tumor progression, our study shows that single-cell resolution, let alone the cells’ spatial configuration, must also be considered for discriminating disease phases and courses.

Analysis of ECM-enriched proteins with progression to tumor was in great part accompanied by corresponding gene expression changes in epithelial and stromal cells. Integration with scRNAseq pinpointed the cellular source down to the granularity of sub-cell types. The increased transcript expression in *M*, *D*, *T* fibroblasts of periostin (POSTN), a developmental ECM protein overexpressed in many tumors^74^, was also increased in the ECM-enriched proteome of *T* samples compared to *mNE* (Fig. 5C). The scRNAseq data (Figs. 3A and 5E) readily attributed its source to the particular subpopulations of fibroblasts that were shared by matched normal gastric samples (*mNS*). Interestingly, this change of expression was partially compensated by a decrease in expression by endothelial cells at more advanced phases of progression (Fig. S7A-B).

Similarly, the marked decrease in expression of some types of collagens in EAC tissues was in accordance with a shift in the subtype of fibroblasts associated with *T* compared to those in *mNE* samples (Fig. 5A). The granular cell-type level resolution of our analyses allowed us to fully appreciate the complexity of such stromal changes, underscoring the importance of the specific cellular source for a change in abundance of a protein in the bulk tissue. Thus, despite the net decrease in collagen-6 protein that could be attributed to a loss of squamous epithelium-associated fibroblasts, this decrease was accompanied by selective increased transcript expression in myofibroblasts (Fig. S7A). This apparent discordance epitomizes the concerted changes in multiple cell types that converge to a pro-tumorigenic tissue landscape. Indeed, whereas COL6-deficient fibroblasts would no longer assemble and maintain a homeostatic ECM, the collagen-6-expressing myofibroblasts would instead drive inflammation and fibrosis via endotrophin, the proteolytic bioactive product of collagen 6^75, 76^ . These COL6-positive myofibroblasts would correspond to the previously described dermoplastic fibroblasts found in melanoma^77^ and BE^15^.

The molecular changes in ECM and cells observed during progression was also manifest in the parallel alteration of mechanical properties of both epithelium and stroma. The reduced epithelium stiffness associated with the switch from squamous to gastric tissue is in agreement with the reported progressive decrease in AFM stiffness in metaplastic CP-A (3.1 kPa) and dysplastic CP-D (2.6 kPa) esophageal epithelial cell lines compared to that of the squamous esophageal epithelial EPC2 cell line (4.7 kPa)^78^, but also in the stroma albeit to a lesser more varied extent. However, progression from *M* to *T* still kept the tissue stiffer than that of inherently softer non-cancerous *mNS* gastric tissue (Fig. 5F), in line with the often observed increase in tissue stiffness in malignancies, which in turn feeds back to modulation of gene expression, including genes that control ECM composition. This mutualism between cellular programs and tissue mechanics underscores the importance of joint analysis of molecular, cellular and mechanical changes^78, 79^ .

Our study also highlighted the importance of considering cell (sub)type abundances as quantitative observables (Fig. 3), illustrated by: (i) the gain of a gastric-specific fibroblast population (Fig. 3A, clusters 2 and 4) at the expense of an esophagus-specific fibroblast population (Fig. 3A, cluster 3) discussed above; (*ii*) a dramatic over-representation of pericyte-like cells (Fig. 3B, clusters 0. 5 and 6) occurring late, i.e. only at the tumor stage; (*iii*) a similar delayed over-representation of VWA1 and PLVAP-expressing endothelial cells (Fig. 3D, cluster 0), two ECM proteins involved in the formation of the stomatal and fenestral diaphragms of blood vessels; (iv) an early and drastic decrease in neutrophils expressing genes of phagocytic and bactericidal function (Fig. 3E, clusters 3 and 7); and (v) a drastic loss of B cell predominance over plasma cells seen in *mNE* but not in *mNS,* and at all phases of progression (*M*, *D*, *T*) (Fig. 3F).

The combined analysis at various size scales of tissue geography enabled by CODEX afforded a new lens through which hitherto unseen tissue organizational changes during progression become visible (Fig. 6P). The identification of consistent cell neighborhoods (NH) and communities (C) of NHs allowed us to quantify changes in the number of cells of particular types (e.g., POSTN^hi^ fibroblasts, CD36^hi^ endothelial cells, p53+ epithelial states) within distinct structures, exposing differentials lost by averaging over large areas. In addition, these structures themselves change in number with progression. For instance, the reduction in neutrophils and concomitant increase in Tregs within the same cell community during EAC progression could not be detected at the global tissue level by scRNAseq. Such changes are of high biological significance, given that the role of Tregs in tissue repair is in part mediated by suppression of neutrophils^80^. The identification of specific multicellular neighborhoods in metaplastic epithelium was consistent with previous descriptions of distinct glandular structures found within BE^70^ .

Shifts in cell state during disease progression from *mNE* to EAC revealed the establishment of an immunosuppressive tumor microenvironment permissive for malignant progression^81, 82^ . Aforementioned POSTN expression, first appearing in *M* fibroblasts has been proposed to serve as a biomarker for BE progression^55^ and is implicated in immunosuppression in the TME^83^ . Furthermore, other highly differentially expressed genes in the POSTN-expressing fibroblast cluster associated with *M*, *D*, *T* included CXCL14, also recently reported in BE scRNAseq analysis^15^ . Notably, CXCL14 has been reported to exert immunosuppressive activities when secreted by fibroblasts but not by epithelial cells^84^ . POSTN and CXCL14 expression may contribute to the local enrichment in Tregs observed in some CODEX cell neighborhoods of *D* and *T* samples. For instance, the “*inflamed mature intestinal”* community which increased in number in *T* samples, contained numerous CD4 Treg cells. Similarly, through consideration of neighborhood structure we found a negative correlation between Tregs and CD36hi endothelial cells in some neighborhoods regardless of the disease phase, in line with the reported association of loss of CD36hi endothelial cells with increased risk of progression in breast cancer^69, 72^ .

At the level of specific genes and pathways associated with EAC progression, numerous inflammatory and malignancy markers in fibroblasts appeared in all modalities: scRNAseq, ECM proteomics and CODEX. Overall, as expected, fibroblasts with gene signatures such as "TGF-beta regulation of extracellular matrix", or “Collagen biosynthesis and modifying enzymes", were enriched with progression as early as metaplasia, in line with adaptive alterations of a stroma subjected to the constant stress of chronic inflammation. Specific changes pointed to a loss of tissue homeostasis involving ECM remodeling by matrix metalloproteinases and collagen chaperones, such as SERPINH1, ultimately resulting in the uncontrolled release of cytokines, such as TGF-beta. Such unopposed TGF-beta signaling might account for the upregulation of periostin expression in fibroblasts at the metaplastic phase^15, 85^ .

Finally, the traditional use of systematic molecular profiling and differential expression/abundance analysis remains potentially useful, notably if we exploit the availability of samples in the progression phases. The identification of a small panel of biomarkers (PRAME, MAGEA6, CASP10, PTPN12, and FAM183A) specifically upregulated at the dysplasia stage (*D*) is of particular practical importance because they could serve as a much sought after biomarker panel for the rare progression from premalignancy (BE) to malignancy (EAC). If confirmed, such biomarkers could initiate and justify more aggressive ablative treatments in any dysplastic BE (including low grade dysplasia) prior to invasive malignancy.

This Atlas of BE progressing to EAC, offers multiple modalities of data that also span multiple size scales from molecular profiles to tissue architecture and mechanics, and will serve as a valuable resource to the research community. Two limitations of this study are: (1) the course of progression is inferred from snapshots of metachronic parallel evolution of lesions as a surrogate of a time course (longitudinal monitoring), which however is permissive given the established sequence of the phases of progression; (2) our matched normal samples used as non-cancerous baseline are actually not disease-free but likely already inflammed tissues as revealed by comparison to disease-free individuals (unpublished observations). Interactive web-portals to interrogate specific molecules in specific cell type and progression phase are available (Methods). Many descriptive but intriguing findings await experimental examination or focused validation in larger cohorts. Clinical translation of some of the observed changes into actionable biomarkers for risk stratification or targets for prevention and intervention holds tremendous potential. It will be also of great interest to determine if our findings can be extended to other chronic inflammation-driven malignancies.

## METHODS

### Sample Collection, Preparation, and Measurements

#### Human Barrett’s Esophagus Tissue Specimens

Fresh tissue specimens were obtained from consented patients with treatment naive Barrett’s esophageal adenocarcinoma (Research Institute - McGill University Health Centre REB # 2007-856) undergoing endoscopy or esophagectomy. They were collected from regions containing tumor, matched normal gastric (gastric cardia) and/or esophageal mucosa (at least 5 cm proximal to the top of columnar lined mucosa), suspected metaplasia and suspected dysplasia. High-Definition white light and narrow band endoscopic imaging was employed to attempt to clinically differentiate dysplasia from non-dysplastic columnar lined mucosa (Barrett’s metaplasia) at the time of surgery. Only tissue specimens with confirmed histological diagnosis were used in subsequent multi-omic analyses (e.g. single cell transcriptomics, multiplex imaging, ECM proteomics). Histological diagnosis was performed on H&E stained sections of formalin fixed paraffin tissue blocks and corroborated by consensus of two expert pathologists (S.C.-B. And P.-O.F.). Collected tissue specimens were divided in equal sections and placed in cold medium (RPMI (Invitrogen) supplemented with Primocin (Invivogen) and gentamycin (Invitrogen)) for single cell RNA sequence processing or shipment to various sites for subsequent analyses. Patient demographics, exposure history (e.g., smoking, proton inhibitor use), Barrett’s extent (Prague Classification), and tumor characteristics (grade, stage) were collected (Suppl. Fig S1A).

### Single-cell RNAseq Methods

#### Single cell dissociation

Tissue specimens were dissected to remove necrotic areas, minced and digested in 5 mL of Advanced DMEM/F12 containing 10 mg Collagenase Type 3 (Worthington) and 500 U Hyaluronidase (Sigma) in a C-tube (Miltenyi) using the gentleMACS Octo Dissociator (Miltenyi). The single cell suspension was resuspended in PBS and 1mM DTT, strained through a 100um cell strainer (Fisher) and spun down (500xg, 5 minutes, 4°C). Cells were resuspended in 0.25% Trypsin-EDTA (Invitrogen) and incubated for 5 minutes at 37°C, followed by addition of 10% fetal bovine serum to inactivate trypsin. The cell pellet (500xg, 5 minutes, 4°C) was resuspended in 2.5U Dispase/10ug DNAse buffer and incubated for 5 minutes at 37°C. The buffer was inactivated by adding excess PBS and the homogenate was strained (40 uM, Fisher) prior to centrifugation (500xg, 5 minutes, 4°C). Red blood cells were lysed using ACK Lysing Buffer (Gibco) for 5 minutes at room temperature, followed by addition of excess PBS, prior to centrifugation (500xg, 5 minutes, 4°C). The cell pellet was finally washed twice with 2% fetal bovine serum in PBS prior to proceeding with single cell capture on the 10x Genomics platform.

#### Single cell suspension quality assessment

Before credentialing the cell suspension, the cells were filtered through a 40 um FLOWMI cell strainer (SP Bel-Art; H13680-0040). Whenever necessary, centrifugation of the cells was carried out at 300xg for 11 minutes. Single cell viability and presence of debris and erythrocytes in the single cell suspension were assessed prior to single cell capturing. Upon adequate viability (i.e. lack of debris and erythrocytes), cells were captured on the 10x Genomics platform.

Cell viability was tested using the ”LIVE/DEAD Viability/Cytotoxicity Kit for mammalian cells” that contained Ethidium Homodimer-1 and Calcein-AM stain (ThermoFisher ; L-3224) dyes. First, a viability stain mix was reconstituted by mixing 0.5 ul of 4mM Calcein-AM, 2ul of 2mM Ethidium Homodimer-1 and 100ul of PBS. A 5ul of cell suspension was then resuspended in 5ul of viability stain mix and the solution was incubated at room temperature for 10 minutes. The sample viability was verified using a hemocytometer (INCYTO C-Chip; DHC-N01-5) through GFP (for the Calcein-AM) and RFP (for the Ethidium Homodimer-1) channels on an EVOS FL Auto Fluorescent microscope (ThermoFisher). Viability was expressed as the percentage of live cells (Calcein-AM / GFP positive cells) over the sum of live (Calcein-AM / GFP positive cells) and dead cells (Ethidium Homodimer-1 / RFP positive) cells.

Erythrocyte contamination was assessed by staining the cells with the cell permeable DNA dye DRAQ5 (ThermoFisher ; 65-0880-92). A nuclear staining mix was made by diluting the DRAQ5 stock solution (5mM) down to 5uM with 1x PBS. Afterwards, 5ul of cell suspension was re-suspended in 5ul of nuclear stain mix and the solution was incubated at room temperature for 5 minutes. The nuclear stain was visualized using a hemocytometer (INCYTO C-Chip; DHC-N01-5) through the Cy5 channel on an EVOS FL Auto Fluorescent microscope (ThermoFisher). Erythrocyte contamination was expressed as the percentage of “round donut-shaped DRAQ5 negative objects on bright-field” over the sum of “round donut-shaped DRAQ5 negative objects on bright-field” and “nuclear stained DRAQ5 positive cells”.

Additionally, we assessed the cell suspension for the presence of any other contaminants/debris as well as contaminants that might interfere with the capturing on the microfluidic chip such as large debris. The percentage of debris presented in the sample was expressed as follows: Percentage of “observed non-cell objects on bright-field” over the sum of “observed objects on bright-field” and “observed cells marked on the fluorescent channels”. A sample was deemed adequate for capturing if “cell viability” was ≥ 70%, “erythrocyte contamination” was ≤ 10% and “debris percentage” was ≤ 30%. The cell concentration in the cell suspension, measured by counting the number of “Calcein-AM / GFP positive cells” and “Ethidium Homodimer-1 / RFP positive cells” in the large 4 squares on each corner of the hemocytometer, was calculated as follows: Number of cells / ul =[(Calcein-AM / GFP positive cells + Ethidium Homodimer-1 / RFP positive cells) / 4] * 10 * 2 where 10 was the dilution factor on the hematocytometer and 2 was the dilution factor when the cell suspension was mixed with the dye solution.

### Single cell capturing

Single cells were captured on the 10x Genomics platform. Single cell 3’ end gene expression profiling was carried out according to the “Chromium Next GEM Single Cell 3’ Reagent Kits v3.1” protocol and recommended reagents. Single cell Copy Number Variation was queried using the “Chromium Single Cell DNA Reagent Kits” protocol and recommended reagents. Of note the CNV kit described above is currently discontinued. The sequencing libraries were created as per the above protocols with the modifications presented in the following section.

### Sequencing of the 10x single cell libraries

Libraries were quantified using a LightCycler 480 Real Time PCR instrument (Roche) and the KAPA library quantification kit (Roche) with triplicate measurements. Library quantification values were used both for the MGI library conversion and for Illumina sequencing normalization.

Libraries sequenced on MGI (MGI Tech) were converted after 10x library construction in order to be compatible with MGI sequencers using the MGIEasy Universal Library Conversion Kit. The kit circularizes the libraries making them compatible for MGI systems. To sequence the circularized libraries, they were first amplified by rolling circle amplification, resulting in a long DNA strand which individually folds into a tight ball (i.e. a DNA nanoball) where one library fragment results in one DNA nanoball. Before loading into the flowcells, the amplified nanoballs were quantified with a Qubit ssDNA HS Assay kit (ThermoFisher), normalized and loaded onto the sequencing flowcell using the auto-loader method (auto-loader MGI-DL-200R). The flowcells have a functionalized surface that captures and immobilizes the nanoballs in a grid pattern. Typically, two libraries were loaded per lane for the single cell RNA libraries. The DNBSEQ-G400RS PE100 MGI kit with App-A primers was used for single cell RNA library sequencing. The DNBSEQ-G400RS PE150 MGI kit with App-A primers was used for single cell DNA library sequencing.

The flowcells were sequenced on a DNBSEQ-G400 MGI sequencer. Single cell RNA libraries were sequenced as follows: 28 cycles for read1, 150 cycles for read2 and 8 cycles for the i7 index. Single cell DNA libraries were sequenced as follows: 151 cycles for Read1, 151 cycles for Read2 and 8 cycles for the i5 index. Because libraries must be color-balanced for all cycles sequenced in order to maintain a minimum ratio of 0.125 for each base at each cycle, color-balanced single index adapters (10x Genomics) were used for libraries sequenced on MGI.

A subset of 23 libraries were sequenced on the Illumina NovaSeq 6000 platform using S4 flowcells. To ensure uniform loading of the libraries, a preliminary pool was sequenced on Illumina iSeq and the library proportions were readjusted accordingly. Another subset of 12 libraries were sequenced on the Illumina HiSeq 4000 system typically with one library per lane.

Although the MGI sequencer has onboard capability to demultiplex samples, we chose to use independent tools to demultiplex the raw fastq files for each lane to give us the flexibility to reprocess if needed. The fastq files generated using the balanced single index adapters were merged for each library after demultiplexing. The MGI runs were mainly demultiplexed by fastq-multx (https://github.com/brwnj/fastq-multx) but also using fgbio/DemuxFastqs (http://fulcrumgenomics.github.io/fgbio/tools/latest/DemuxFastqs.html). In both instances we used a mismatch of 1. Illumina runs were demultiplexed using the the standard bcl2fastq tool.

### scRNA-seq Data Processing and Analysis

#### Read processing and alignment

After polyA-trimming via cutadapt (v3.2)^86^, reads were pseudo-aligned to the GRCh38 reference transcriptome (ENSEMBL release 96) with kallisto (v0.46.2)^87^ using the default kmer size of 31. The pseudo-aligned reads were processed into a cell-by-gene count matrix using bustools (0.40.0)^88^ . Cell barcodes were filtered using the whitelist (v3) provided by 10xGenomics. All further processing was done in scanpy (v.1.7.1)^89^.

#### Quality control and normalization

Quality control was performed for each sample independently as follows. Cell barcodes with less than 1000 counts or less than 500 genes expressed or with more than 10% mitochondrial gene expression were removed. Doublet cells were identified using scrublet^90^, and any cell barcode with a scrublet score > 0.2 was removed. Only coding genes were retained in the final count matrix. Expression profiles were normalized by total counts, the 4000 most highly variable genes being identified ^91^, renormalized, log-transformed and z-scored. The data were projected onto the first 50 principal components.

After the above per-sample preprocessing, samples were pooled and integrated using Harmony^20^ on the first 50 principal components with a maximum of 25 iterations. A nearest neighbor graph (k=15) was calculated on the Harmony-corrected principal components space. Datasets were visualized in 2D via UMAP^92^ and initialized with PAGA^93^ coordinates. The nearest neighbor graph was clustered with the Leiden algorithm^94^ . The processed datasets were visualized interactively using Cellbrowser^95^ allowing for easy access and exploration across teams and laboratories.

#### Cell type calling

Cell types were called using the Human Cell Landscape (HCL) reference dataset ^21^ . Briefly, the raw expression profile *x_i_* of cells *i* was normalized by total counts and log-transformed 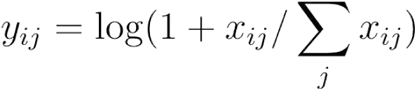 with *x_ij_* the counts of genes *j* in each cell *i*. Cells were compared to the Human Cell Landscape reference by Pearson correlation, and the reference profile with the highest correlation determined the cell type call. If the highest correlation was below 0.3, the cell type was defined as "unknown".

The per-cell types were categorized on a per-cluster level based priors calls with the expression of markers indicated from a range of sources including Cell Ontology^96^ and reference materials from Thermofisher, Abcam and BioLegend.

Fine-grained labels were produced using a number of techniques. To fine grain the squamous esophageal epithelium we compared our esophageal dataset to 6 esophageal samples from the Human Cell Atlas^97^ . HCA-derived gene signatures were produced from the top 50 most differentially expressed genes for each cell type from the HCA data. Using the single-cell gene set scoring method GSSNNG, we scored the cell type labels using those HCA-derived signatures. The squamous epithelium subtypes (upper, intermediate, basal) were clearly indicated through high gene set scores in the CRUK dataset. Also, the HCA signatures clearly identified our cell types including fibroblasts and glandular gastro-intestinal (GI) epithelium. Some HCA signatures were non-specific in our dataset including the esophageal basal cell layer, the venous endothelium, and the ductal epithelium. Otherwise, fine graining was performed by using well known marker genes from the literature.

#### Sample PCA and clustering

In order to apply PCA, sample level clustering, and perform differential expression (as described below), pseudo-bulk profiles were produced for each coarse-grained cell label and over all cells. To produce a pseudo-bulk profile computationally, cells of a given type were selected and raw gene expression counts were summed. For a given cell type, such as for example CD8 T cells, this produced one gene expression profile per sample. PCA was performed using R ’prcomp’, principal components (eigenvectors) were recovered and used for plotting of principal components. Clustering of samples was performed using the R hclust function with average linkage.

#### Differential expression

In order to estimate differential expression between the tissue diagnosis on a per cell basis, we applied DESeq2 to pseudo-bulk profiles while accounting for sequencing depth and patient heterogeneity using the model *gene ∼ patient + cell_counts + avg_molecules + dx* where the *patient* is patient ID (each patient has several different samples), *cell_counts* is the number of cells observed in a given sample, *avg_molecules* is the mean of gene counts, *dx* is the tissue diagnosis. The goal is to find genes where the variations in expression patterns are explained more by the change in tissue diagnosis rather than from other considered factors.

There was considerable bias introduced from ambient RNA which produced false positive results^98^ . In order to identify differentially expressed genes that were artifacts the following heuristic was used: If a gene was found to be differentially expressed in a cell type and was also differentially expressed in other cell types, and the ambient profile was correlated with the pseudo-bulk profile (across samples) and statistically associated with the tissue diagnosis [other metrics], then the DEG was removed from the results. Gene expression heatmaps were created by taking the collection of differentially expressed genes, and scoring genes based on association with disease progression. In particular, for the purpose of visualization, genes were selected through association with (*mNE*, *mNS*, *M*) vs. (*D*,*T*) using logistic regression.

#### Gene set scoring

For each coarse-grained cell type, differentially expressed genes with a max log2 fold change (between tissue diagnoses) of at least 0.58 and adjusted p-value of 0.05 were submitted to the Enrichr service^26^. Pathway enrichment tables were downloaded for BioPlanet 2019, MSigDB Hallmarks, KEGG 2021, and WikiPathways 2021. (see Suppl. Table S2)

#### Statistical analysis of cell type proportion changes

Changes in cell type proportions across diagnoses were analyzed with scCODA^99^ . Changes were computed relative to mNE samples and reference cell types were selected based on the cell type with the least dispersion across diagnosis. To account for inter-patient differences, and systematic differences in cell type composition between biopsy/resection samples, those were included as covariates into the scCODA model. Model inference was performed using HMC and 60000 iterations. MCMC chains were inspected manually for convergence. Statistically significant covariates were determined using the model’s posterior inclusion probabilities and an FDR of 0.05.

#### Copy Number Variation inference from scRNAseq

We used a python re-implementation^100^ of the inferCNV algorithm ^48^ to call copy number variation (CNV) in the single cell data. First, all samples across all diagnoses were pooled and epithelial cells were extracted based on cell type calling. Any epithelial cell from a mNE/mNS sample was assigned to the reference set of inferCNV, any other epithelial cell was assigned to the query set in inferCNV. After filtering out lowly expressed genes (mean expression <0.1) and standard inferCNV preprocessing, the data were smoothed along chromosomal coordinates with a window size of 101 and a step size of 2. The CNV burden $s_i$ of a single cell $i$ was estimated as $s_i = \sum_j |X_{ij}|$, where $X_{ij}$ is the inferred gain/loss in cell $i$, gene window $j$. The 99% quantile of the CNV burden $s_i$ in mNE/mNS epithelial cells was calculated and, assuming no CNV’s present in mNE/mNS samples, any non-reference cells exceeding this threshold were classified as containing CNVs.

### Proteomics Methods

#### Chemicals

LC-MS-grade acetonitrile (ACN) and water were obtained from Burdick & Jackson (Muskegon, MI). Reagents for protein chemistry, including sodium dodecyl sulfate (SDS), ammonium bicarbonate, iodoacetamide (IAA), dithiothreitol (DTT), sequencing-grade endoproteinase Lys-C, and formic acid (FA) were purchased from Sigma-Aldrich (St. Louis, MO). Sequencing-grade trypsin was purchased from Promega (Madison, WI). Glycerol-free PNGase F was purchased from New England BioLabs (Ipswich, MA).

#### Sample Preparation

Proteomic analysis was performed as described in Bons *et al.*^101^ . Briefly, fresh esophageal tissues were minced in small pieces, weighed, and flash frozen for storage at -80⁰C. The ECM fraction was isolated from the frozen tissues using the Compartment Protein Extraction Kit (Millipore, #2145) as per manufacturer’s protocol. About 1/10 of purified ECM was used to assess ECM protein enrichment purity and yield by Western blot analysis. From the remaining purified ECM fraction, proteins were solubilized by agitation for 10 minutes in a solution containing 1% SDS, 50 mM DTT and 1X NuPAGE lithium dodecyl sulfate (LDS) sample buffer (Life Technologies, Carlsbad, CA), followed by sonication for 10 minutes, and finally heating at 85 ⁰C for 1 hour with agitation. Solubilized proteins were concentrated in a single stacking acrylamide Bis-Tris gel, in-gel reduced with 10 mM DTT, and alkylated with 55 mM IAA. Finally, proteins were in-gel digested with 250 ng of sequencing-grade endoproteinase Lys-C in 25 mM ammonium bicarbonate at 37 ⁰C for 2 hours with agitation, followed by an overnight incubation with 250 ng sequencing-grade trypsin in 25 mM ammonium bicarbonate at 37 ⁰C with agitation. After tryptic peptide extraction, samples were vacuum dried, resuspended in 25 mM ammonium bicarbonate, and peptides were deglycosylated with 3 µL (1,500 U) of glycerol-free PNGase F at 37 ⁰C for 3 hours with agitation. The reaction was quenched by adding FA to a final concentration of 1%. Peptide samples were finally desalted using stage-tips made in-house containing a C_18_ disk, vacuum dried, and re-suspended in aqueous 0.2% FA spiked with indexed retention time peptide standards (iRT, Biognosys, Schlieren, Switzerland)^102^ .

#### LC-DIA-MS Analysis

LC-MS/MS were performed on an Eksigent Ultra Plus nano-LC 2D HPLC system (Dublin, CA) combined with a cHiPLC system directly connected to an orthogonal quadrupole time-of-flight (Q-TOF) SCIEX TripleTOF 6600 mass spectrometer (SCIEX, Redwood City, CA). The solvent system consisted of 2% ACN, 0.1% FA in H_2_O (solvent A) and 98% ACN, 0.1% FA in H_2_O (solvent B). Proteolytic peptides were loaded onto a C_18_ pre-column chip (200 μm × 6 mm ChromXP C18-CL chip, 3 μm, 300 Å; SCIEX) and washed at 2 μL/minute for 10 minutes with the loading solvent (0.1% FA in H_2_O) for desalting. Peptides were transferred to the 75 μm × 15 cm ChromXP C_18_-CL chip, 3 μm, 300 Å (SCIEX) and eluted at 300 nL/min with the following gradient of solvent B: 5% for 5 min, linear from 5% to 8% in 15 min, linear from 8% to 35% in 97 min, and up to 80% in 20 min, with a total gradient length of 180 min. Samples were analyzed by data-independent acquisition (DIA) using 64 variable-sized windows covering the m/z 400-1,250 range (Suppl. Table S7)^103–105^ . MS scans were collected with 250-ms accumulation time, and MS/MS scans with 45-ms accumulation time in “high-sensitivity” mode. The collision energy (CE) for each segment was based on the z=2+ precursor ion centered within the window with a CE spread of 10 or 15 eV.

#### MS DIA Data Processing

DIA data were processed in Spectronaut (version 14.10.201222.47784) (Biognosys) using a pan-human library containing 10,316 proteins^106^ . Data extraction parameters were selected as dynamic using non-linear iRT calibration. Identification was performed using 1% precursor and protein q-values. Quantification was based on the MS/MS peak areas of the 3-6 best fragment ions per precursor ion, local normalization was applied, and iRT profiling was selected. Differential protein abundance analysis was performed using paired t-tests, and p-values were corrected for multiple testing using the Storey method^107^ . Protein groups with at least two unique peptides, q-value ≤ 0.001, and absolute Log2(fold-change) ≥ 0.58 were considered to be significantly altered (Suppl. Table S6).

#### Atomic Force Microscopy Methods

Snap frozen patient samples were cryosectioned. A reference slide has been H&E stained from the adjacent test sample to visualize the location of epithelium and stroma. Unfixed slides were tested with AFM for stiffness and distribution of measurements were visualized using AFM manufacturer’s software (Fig. S8).

### CODEX Multiplexed Protein Imaging Methods

#### CODEX Array Creation and Pathology Annotation

Imaging data were collected from 5 human donors, each of whom constituting a dataset. Each dataset included tissue sections taken from individually diagnosed formalin fixed paraffin embedded (FFPE) tissue blocks that were combined onto the same coverslip (cut at 4 µm thickness). To ensure accurate disease phase diagnosis, three pathologists independently evaluated the H&E staining of the sections performed on the same tissue sections as used for the CODEX multiplexed imaging (Fig. S9B-C). They called disease phase granular diagnosis (e.g., mNE, mNS, M), and estimated percentages of type of epithelium in each image (e.g., % squamous, % metaplasia, % dysplasia, % tumor). The pathologists scores were then aggregated and averaged (Suppl. Table S8).

#### CODEX Antibody Conjugation and Panel Creation

CODEX multiplexed protein imaging was executed according to the CODEX staining and imaging protocol previously described^63^. Antibody panels were chosen to include targets that identify subtypes of intestinal epithelium and stromal cells, and cells of the innate and adaptive immune system. Detailed panel information can be found in Suppl. Table S9. Each antibody was conjugated to a unique oligonucleotide barcode, after which the tissues were stained with the antibody-oligonucleotide conjugates and validated to ensure that staining patterns matched expected patterns already established for IHC within positive control tissues of the esophagus or tonsil. Similarly, Hematoxylin and Eosin morphology stainings were used to confirm location of marker staining. First, antibody-oligonucleotide conjugates were tested in low-plex fluorescence assays and signal-to-noise ratio was also evaluated at this step, then conjugates were tested all together in a single CODEX multicycle.

#### CODEX Multiplexed Protein Imaging

The tissue arrays were then stained with the complete validated panel of CODEX antibodies and imaged^63^ . Briefly, the workflow entailed cyclic stripping, annealing, and imaging of fluorescently labeled oligonucleotides complementary to the oligonucleotide on the conjugate. After validation of the antibody-oligonucleotide conjugate panel, a test CODEX multiplexed assay was run, during which signal-to-noise ratio was again evaluated, and the optimal dilution, exposure time, and appropriate imaging cycle was evaluated for each conjugate (Suppl. Table S9). Finally, each coverslip array underwent CODEX multiplexed imaging.

#### CODEX Data Processing

Raw imaging data were then processed using the CODEX Uploader for image stitching, drift compensation, deconvolution, and cycle concatenation. Processed data were then segmented using the CellVisionSegmenter, a neural network R-CNN-based single-cell segmentation algorithm (https://github.com/michaellee1/CellSeg) ^108^. After upload, the images were again evaluated for specific signal: any marker that produced an untenable pattern or a low signal-to-noise ratio was excluded from the ensuing analysis. Uploaded images were visualized in ImageJ (https://imagej.nih.gov/ij/).

#### CODEX Cell Type Analysis

Cell type identification was done following the methods developed previously ^67^ . Briefly, nucleated cells were selected by gating DRAQ5, Hoechst double-positive cells, followed by z-normalization of protein markers used for clustering (some phenotypic markers were not used in the unsupervised clustering). The data were overclustered with X-shift (https://github.com/nolanlab/vortex). Clusters were assigned a cell type based on average cluster protein expression and location within the image. Impure clusters were split or reclustered following mapping back to the original fluorescent images.

#### CODEX Cell-cell colocalization analysis and Shannon’s Index

To evaluate enriched cell-cell interactions, we calculated the frequency of neighbors using a nearest neighbor (n=10 neighbors) approach and compared the frequency of occurrences to 10,000 permutations of the cell type locations. We filtered this list for cell-cell interactions enriched within certain conditions compared to the other disease states. Shannon’s Diversity Index was calculated by taking the negative sum of each proportion multiplied by the natural logarithm of the proportion.

#### CODEX Neighborhood and Community Identification Analysis

Neighborhood analysis was performed as described previously^109, 110^. Briefly, this analysis involved (i) taking windows of cells across the entire cell type map of a tissue with each cell as the center of a window, (ii) calculating the number of each cell type within this window, (iii) clustering these vectors, and (iv) assigning overall structure based on the average composition of the cluster. Neighborhoods were overclustered to 30 clusters. These clusters were mapped back to the tissue and evaluated for cell type enrichments to determine overall structure and merged down into the final unique neighborhoods. Communities were determined similar to how multicellular neighborhoods were determined with some minor differences ^65^ . Briefly, the cells in the neighborhood tissue maps were taken with a larger window size of 100 nearest neighbors. These windows were then taken across the entirety of the tissue and the vectors clustered with k-means clustering and overclustering with 20 total clusters. These clusters were mapped back to the tissue and evaluated for neighborhood composition and enrichment to determine overall community type.

## Supplemental Figures

**Figure S1.**
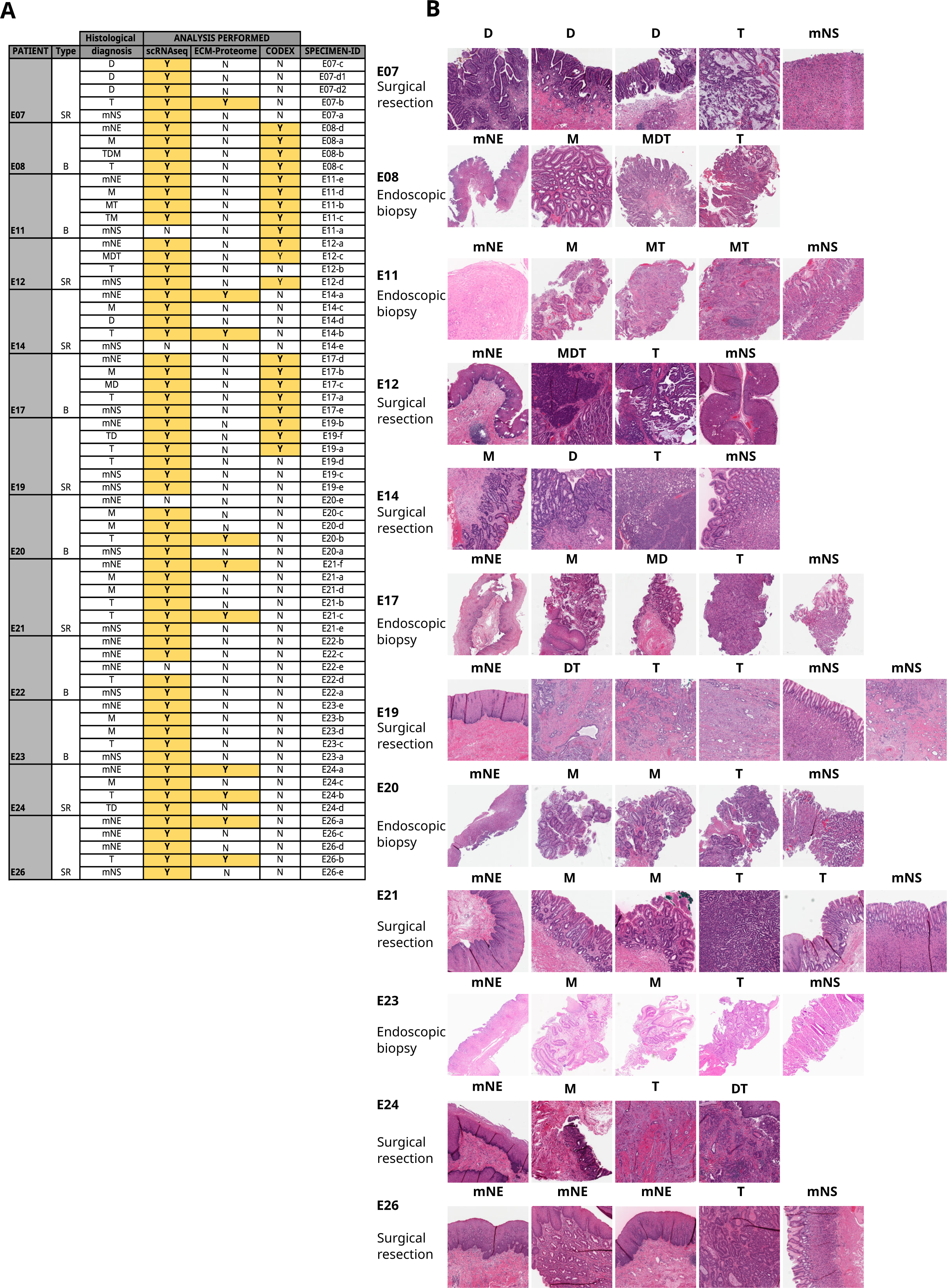
Description of clinical samples and multi-omics analyses conducted. **(A)** Table describing the samples analyzed in this study including sampling method (surgical resection (SR) or endoscopic biopsy (B)), tissue diagnosis (*mNE*: matched Normal Esophagus, *M*: Barrett’s metaplasia, *D*: dysplasia, *T*: tumor, *mNS*: matched Normal Stomach - other abbreviations indicate a mixture of tissue types), type of analysis performed and specimen ID. **(A) (B)** Selection of pathohistological images of scanned hematoxylin and eosin-stained tissue sections independently assessed by two expert esophageal pathologists to confirm tissue diagnosis.

**Figure S2.**
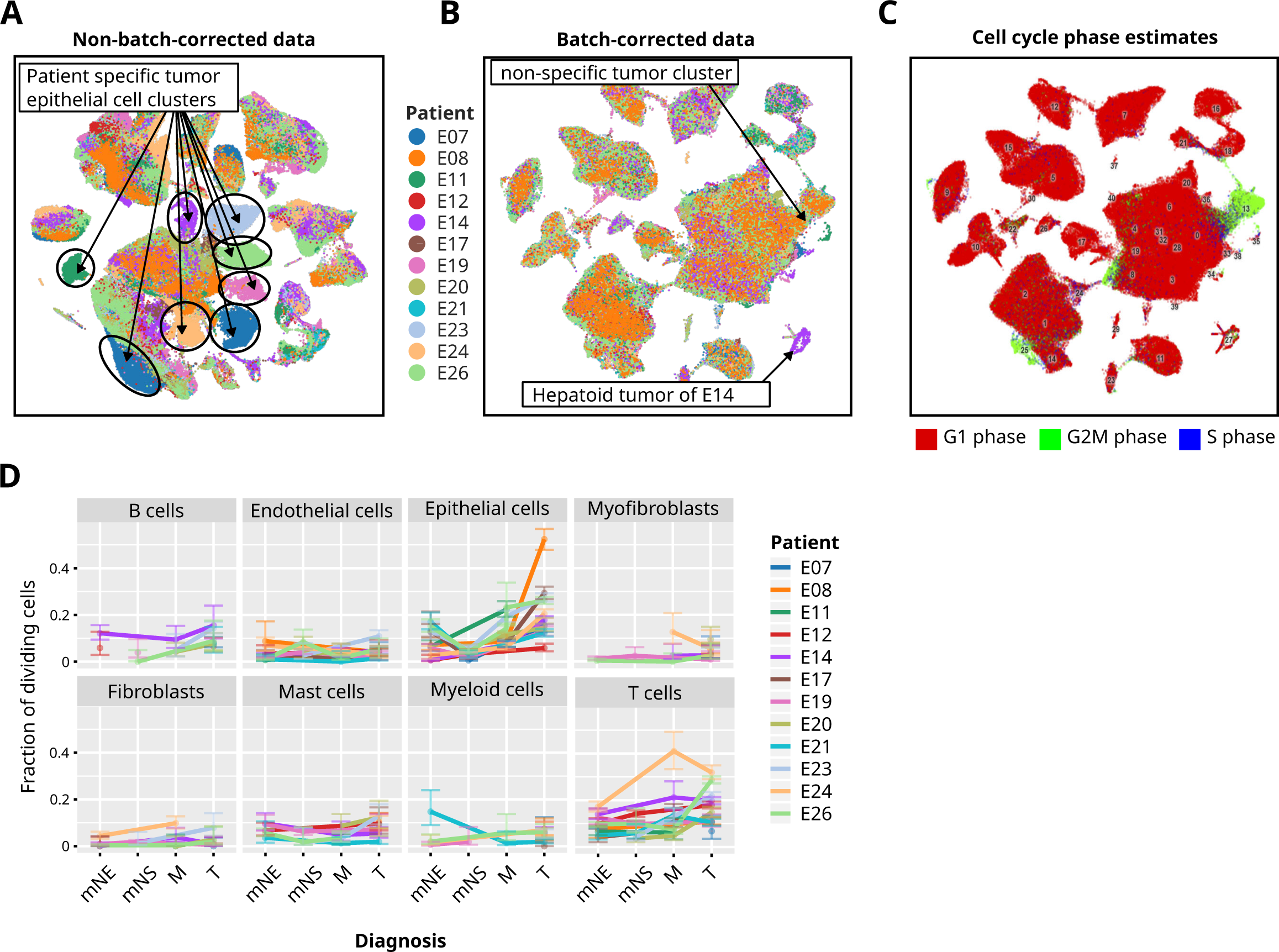
Overview of sample space and global cell type survey. **(A)** UMAP visualization for non-batch corrected scRNAseq data of all analyzed cells across all histological diagnoses (progression phases), color-coded by cluster. Note that the non-batch corrected data show separation into patient-specific cancer cells. **(B)** UMAP visualization for batch corrected scRNAseq data of all analyzed cells across all histological diagnoses, color-coded by cluster. **(A) (C)** Cells color-coded for predicted cell cycle phase (Methods). Subsets of epithelial tumor cells and T cells appear highly proliferative (G2M phase, green). **(D)** Fraction of proliferating cells (S or G2M phase) color-coded for each patient for each cell type and phase of progression (diagnosis). Error bars denote 5% and 95% Bayesian credibility intervals of a binomial model.

**Figure S3.**
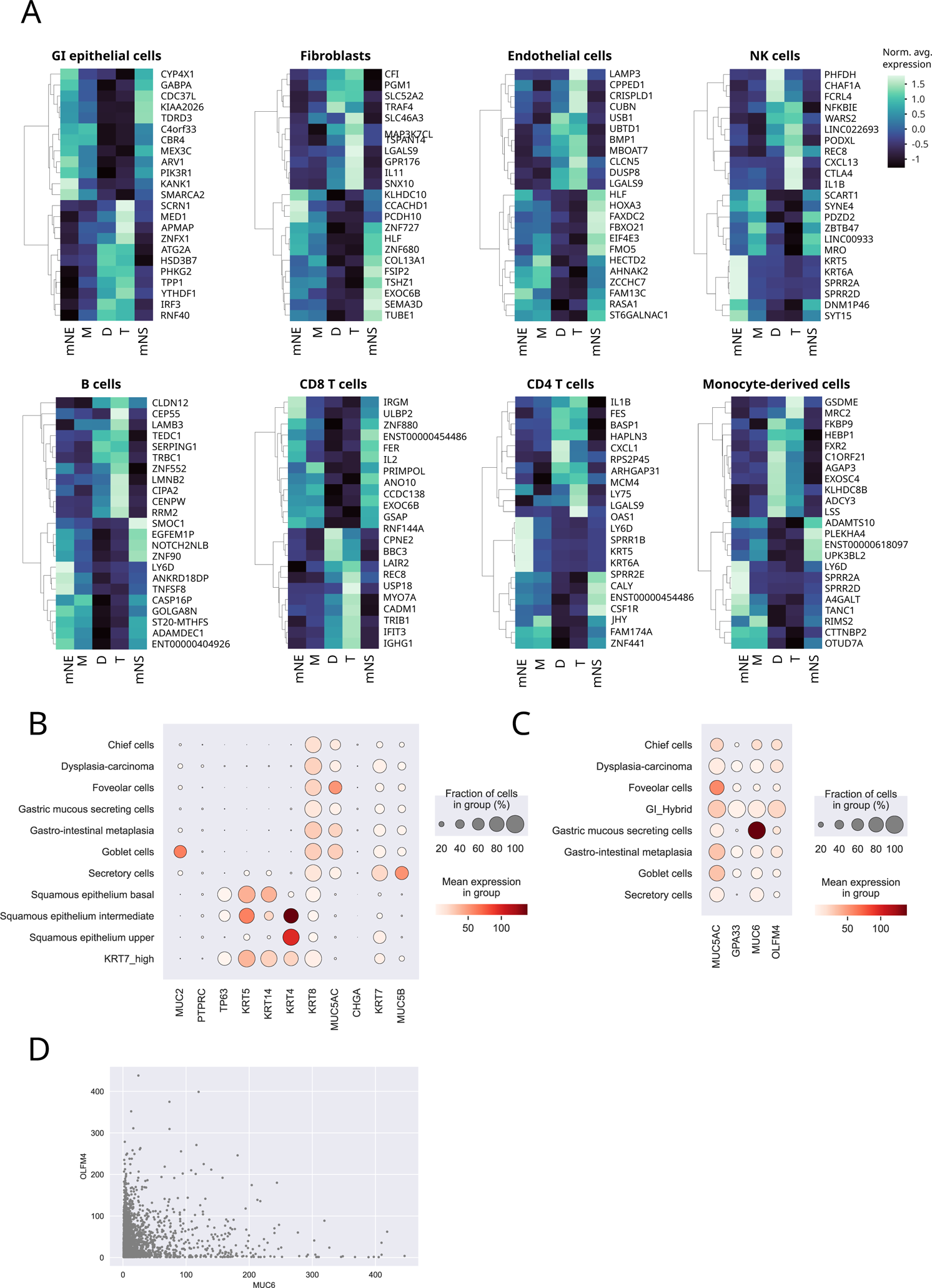
Cell type-specific signatures of gene expression changes during BE progression. Differential gene expression at each phase of progression (diagnosis) based on pseudo-bulk transcriptomes, for 8 major cell types. The observed gene expression patterns specific for various cellular compartments illustrate profound epithelial, stromal and immune tissue alterations associated with disease progression.

**Figure S4.**
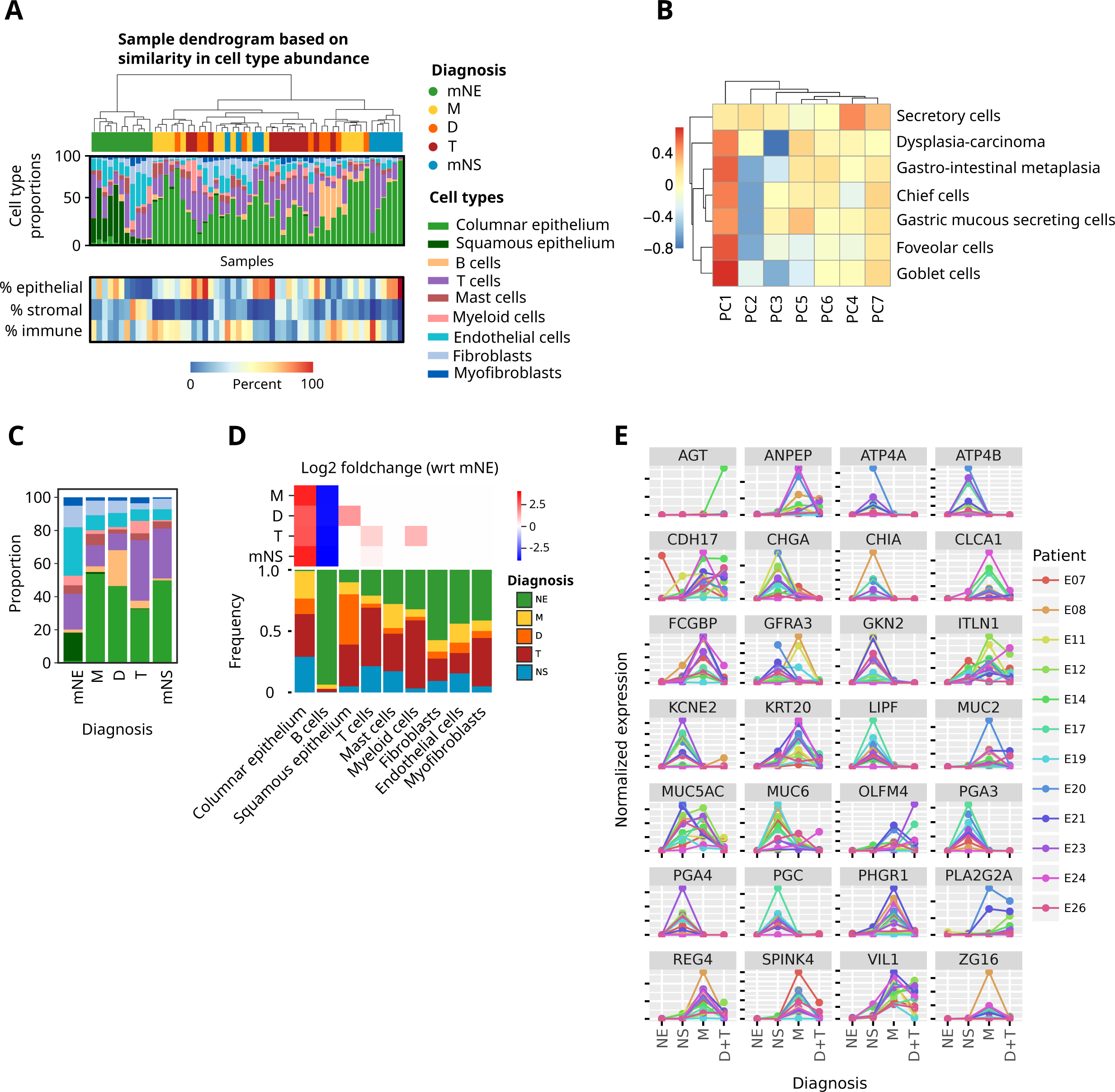
Cell type composition changes during BE progression. **(A)** TOP: Clustering of individual samples based on cell type composition (proportions) Each sample’s diagnosis is indicated above the dendrogram. BOTTOM: Epithelial, stromal and immune percentages of each sample (aggregated from cell type proportions used for clustering) are shown in the heatmap below the dendrogram. **(B)** Sample level PCA performed with pseudo-bulk gene expression measures showed that the variation in the PCA was largely due to variation in cell type quantities as shown by correlation with the principal coordinates and cell type proportions across samples. **(C)** Cell type proportions (as shown in A) collapsed by diagnosis. **(D)** Significant changes in cell type composition across diagnoses. Statistically significant changes with respect to *mNE* (determined by scCODA, see Methods) are shown above the bar graph (red: upregulated, blue: down-regulated). **(E)** Normalized expression of differentiation markers in epithelial cells across the different phases of disease progression shows a loss of differentiation at the D/T phase of progression.

**Figure S5:**
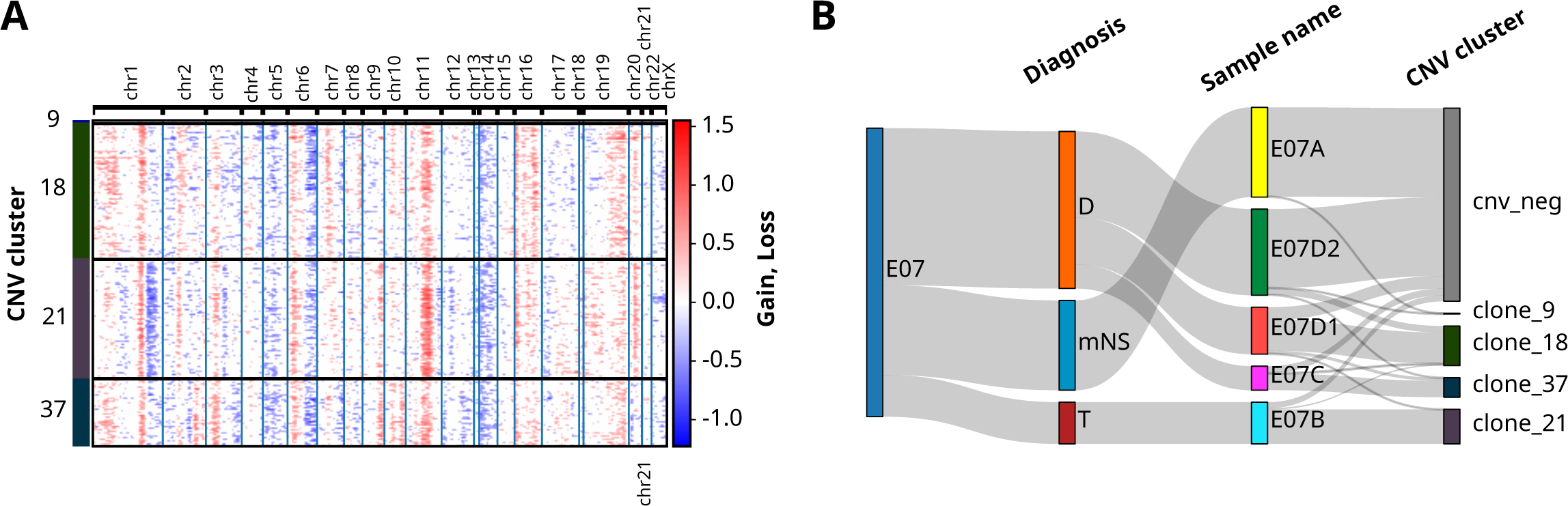
Copy number profile (A) and distribution of copy number clusters/clones across samples and diagnoses (B) inferred from scRNAseq. While dysplastic (*D*) cells (samples E07C and E07D1) form their own CNV clusters (clones 18 and 37), cells from matched *T* (sample E07B) show similar CNVs (clone 21), e.g., a gain of chr6p and a loss of chr6q indicating a clonal relationship between *D* and *T*.

**Figure S6.**
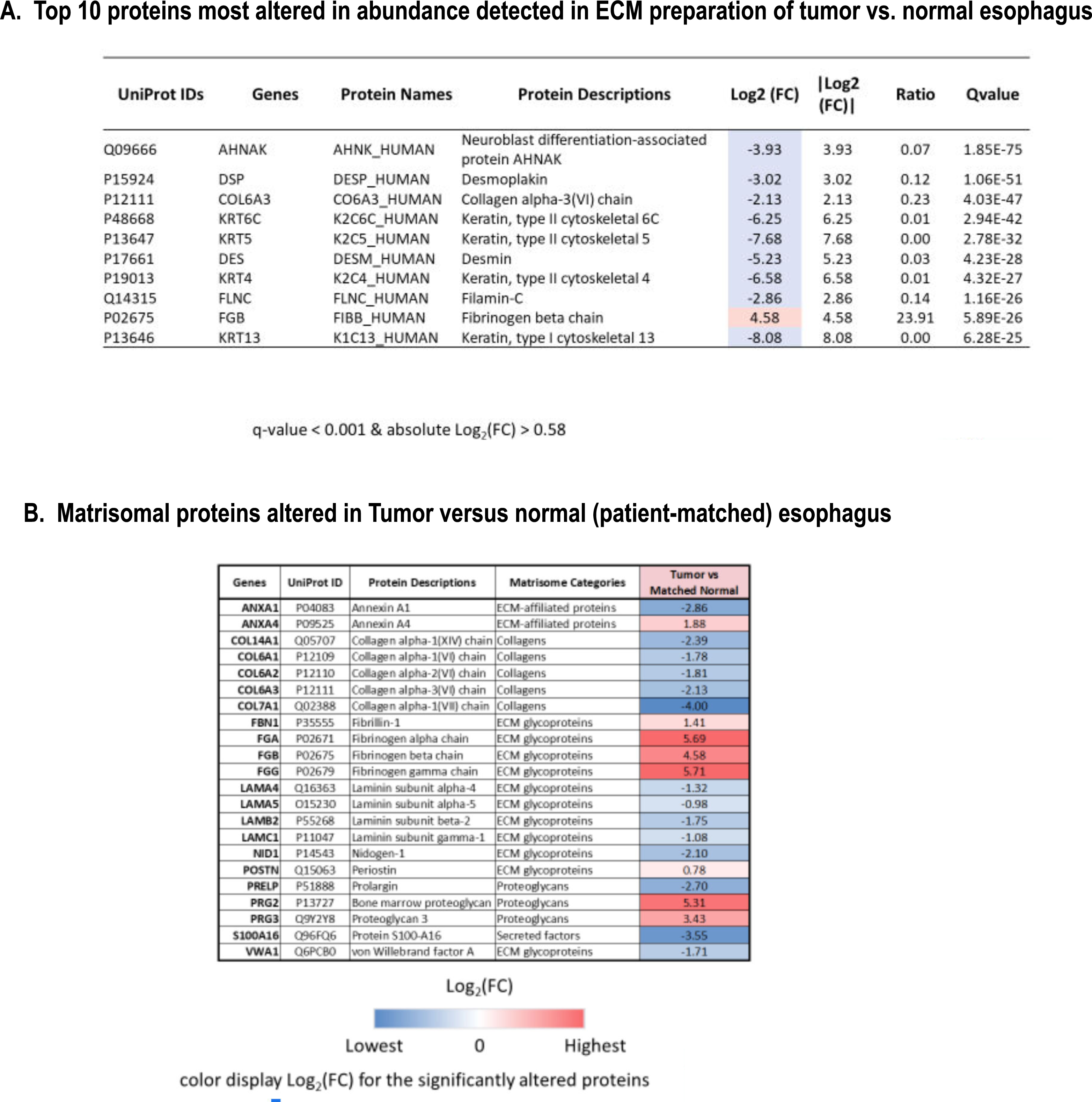
Protein abundance changes in EAC (*T*) compared to matched normal esophagus (*mNE*). **(A)** Top 10 proteins with most altered abundance detected in ECM-enriched samples or *T* vs. *mNE* tissues. **(B)** Core matrisomal protein signatures altered in *T* versus *mNE*.

**Figure S7.**
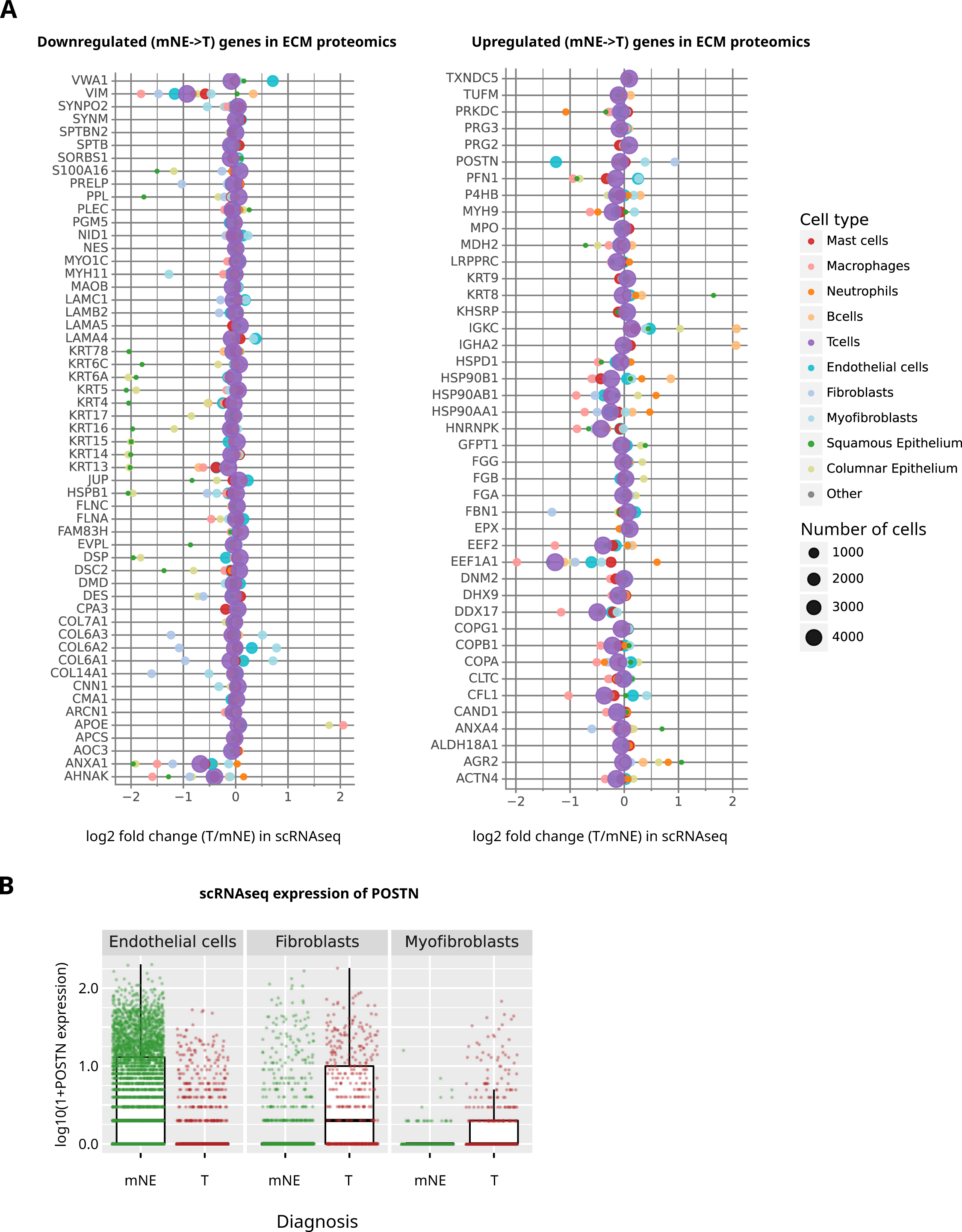
(A) Identification of likely cell source of proteins with altered abundances during BE progression. Gene expression fold changes in scRNAseq pseudo-bulk analysis were queried for proteins that were identified as differentially abundant in the ECM proteomic analysis, i.e., either decreased (left panel) or increased (right panel) in *T* vs. *mNE*, respectively (Fig. 5 and Table S5). Cell type specificity and abundance for each transcript is displayed as circles (color-coded for cell type and size-scaled for cell abundance). **(B)** Plots with changes in mRNA expression of periostin (POSTN) inferred from scRNAseq data, grouped by cell type and diagnosis, reveals complexity changes that depends on cell source (shown in A).

**Figure S8.**
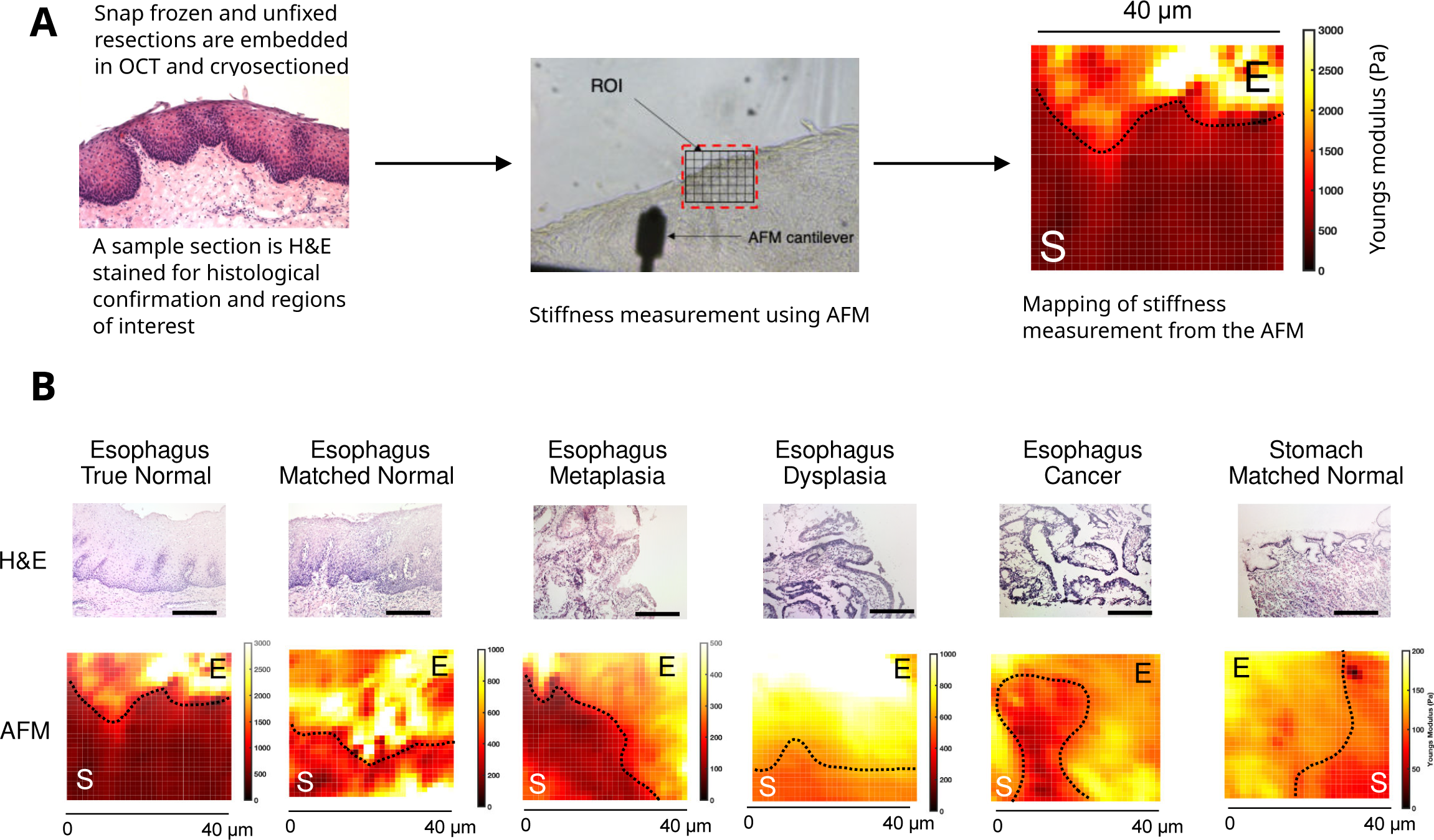
AFM mechanics is consistent with a shift in tissue identity from *squamous esophagus* to *BE* (esophagus to stomach). **(A)** Workflow of sample preparation for AFM stiffness measurements. **(B)** Stromal and epithelial esophagus stiffness gradients were compared at various phases of progression to EAC. Representative H&E images and corresponding AFM stiffness maps showing difference between stroma and epithelium stiffness at different phases of esophageal cancer progression and comparison with matched normal squamous esophagus (i.e., *mNE* adjacent to *BE* and tumor) and true normal (disease-free) squamous esophagus tissues. Dashed lines indicate the interface between stroma (S) and epithelium (E). Scales: 200 μm.

**Figure S9.**
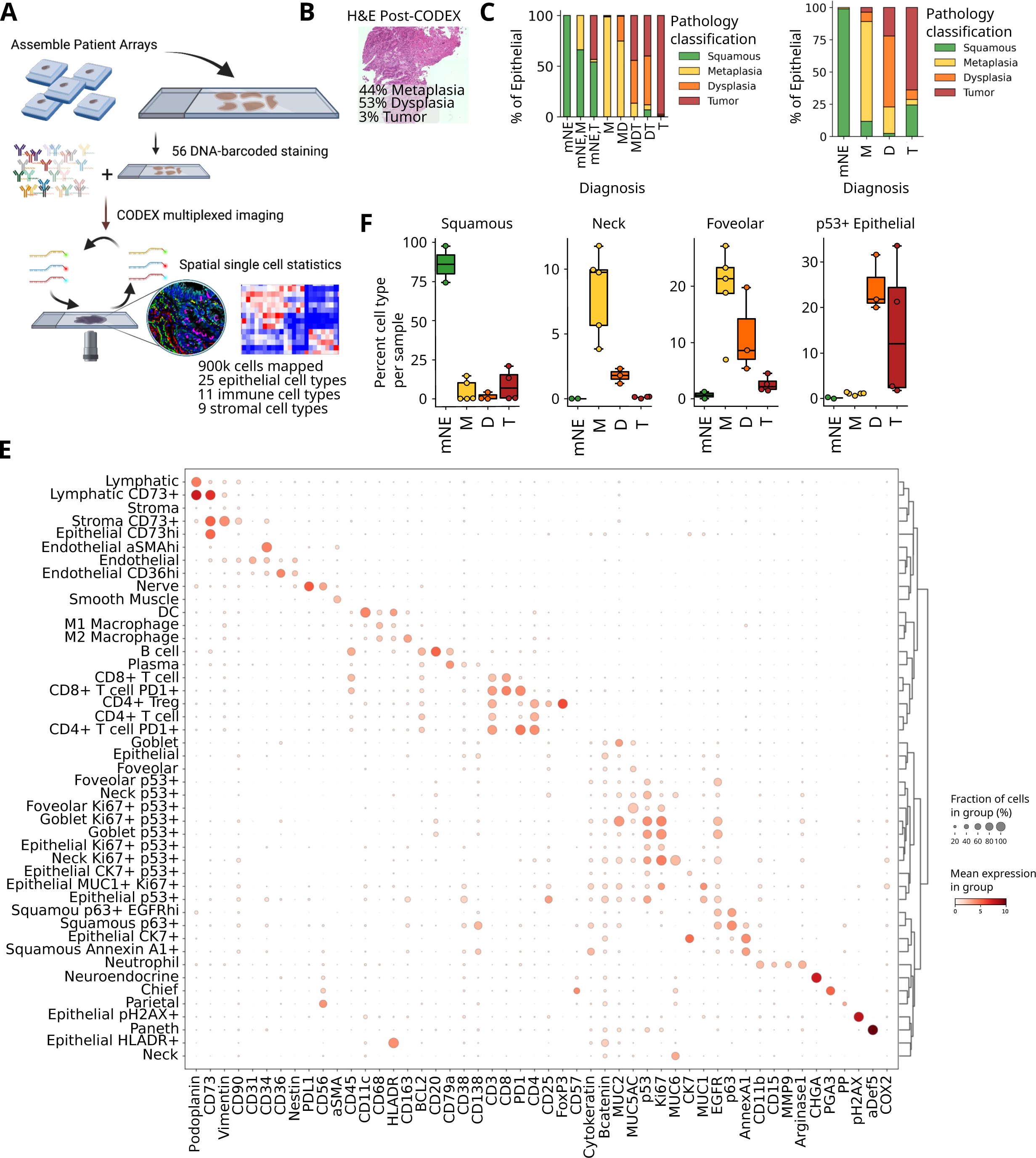
Cell type-specific analysis from CODEX multiplexed proteomics imaging of samples corresponding to disease progression phases of BE-associated EAC. **(A)** Schematic for overall approach to image tissues with CODEX imaging. **(B)** Representative H&E image taken after CODEX imaging with percentage of diagnosis (based on epithelial cell histology) labeled by pathologists. **(C-D)** Quantification of epithelial percentage (*y*-axis) for each overall (C) sub and (D) major diagnosis by pathologist from the H&E stain of the tissue section following CODEX multiplexed imaging. **(E)** Average protein expression (*x*-axis) by cell type (*y*-axis) derived from the clustered CODEX multiplexed imaging data. **(F)** Epithelial cell type percentages across the different disease phases as diagnosed by pathologists.

**Figure S10.**
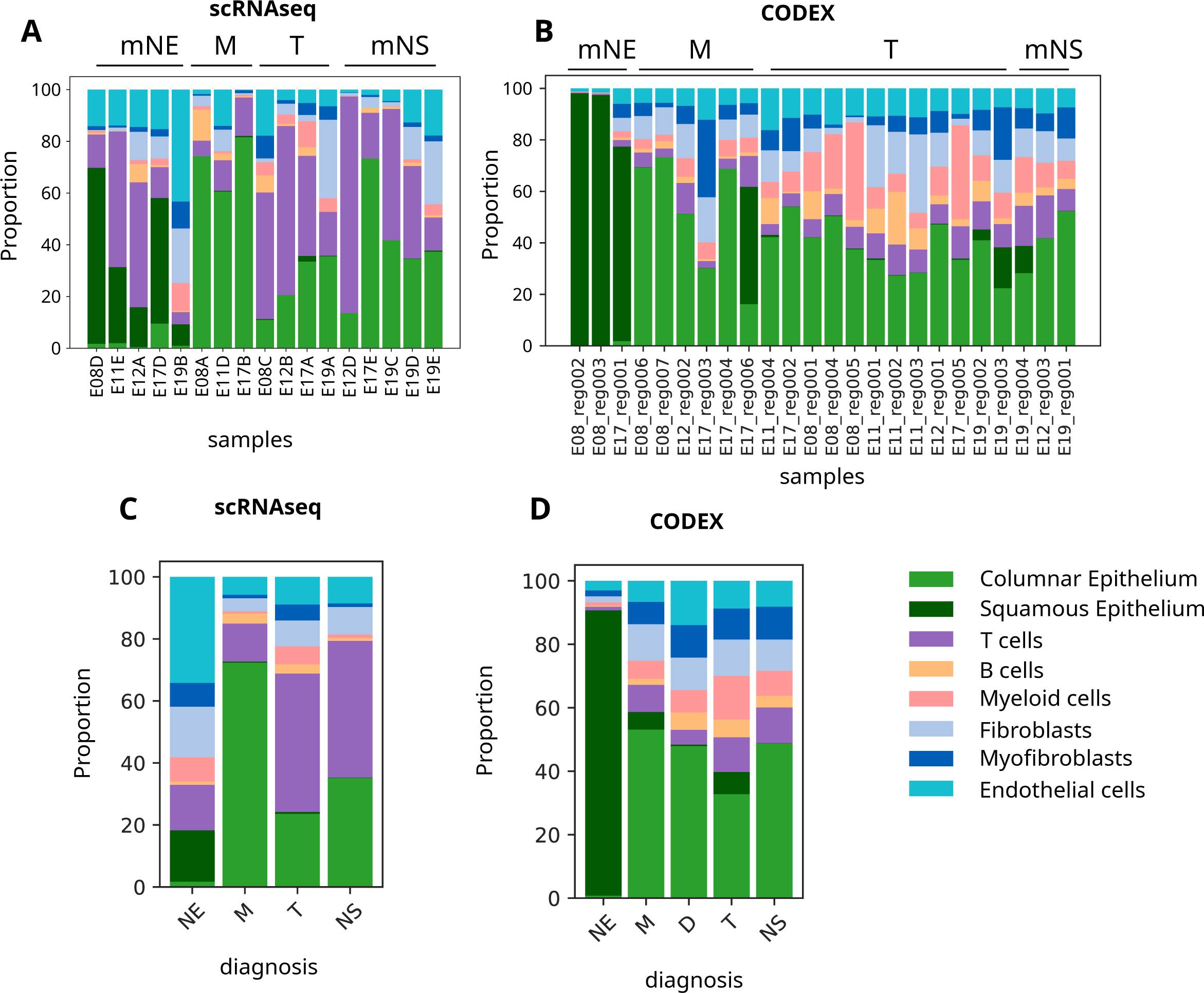
Comparison of cell type abundances across patients analyzed by both scRNAseq (A, C) (clinical samples) and CODEX (B, D) (adjacent tissue regions) grouped by sample (A, B) and diagnosis (C, D).

**Figure S11.**
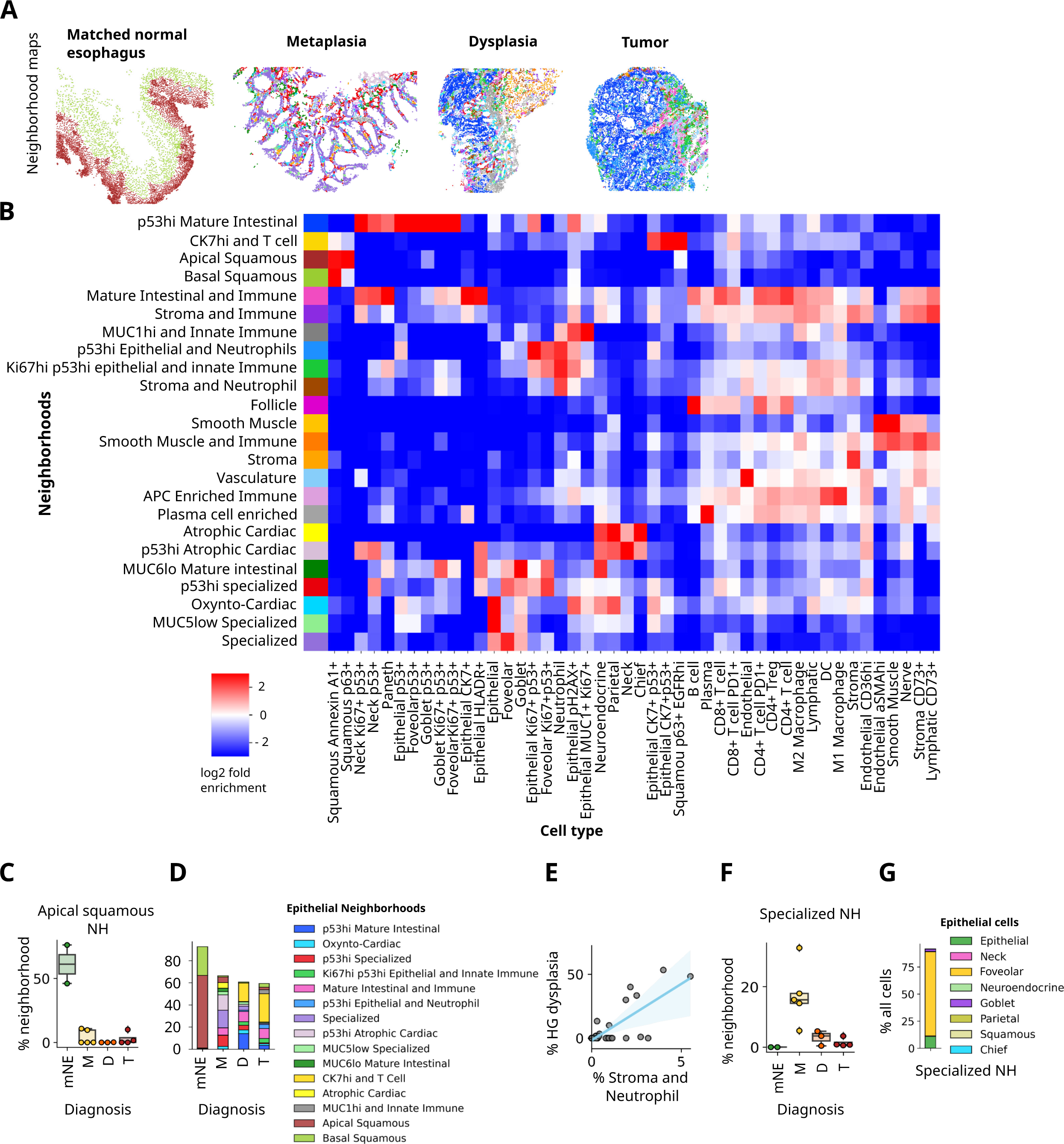
Multicellular Neighborhood (NH) analysis by CODEX imaging reveals multicellular neighborhood tissue reorganization during the progression of BE to EAC. **(A)** Maps of multicellular neighborhoods mapped back to original coordinates for one representative patient (5 patients total). **(B)** Definition of NHs based on cell types enrichment (red) or depletion (blue) within individual neighborhoods identified in Figure S8E. Rows represent hence -identified NHs, and their name given on the left. **(C)** Percentage of “Apical Squamous” NH within the different progression phases. **(D)** Average percentage of the epithelial neighborhoods within each of the phases as determined by a pathologist post H&E staining. **(E)** Percentage of high-grade (HG) dysplastic epithelium as determined by a pathologist (*y*-axis) versus percentage of “Stroma and Neutrophil” NH determined by CODEX for each of the 27 regions imaged. **(F)** Percentage of “Specialized” NH within the different disease phases. **(G)** Percentage of epithelial cell types within the “Specialized” NH.

**Figure S12.**
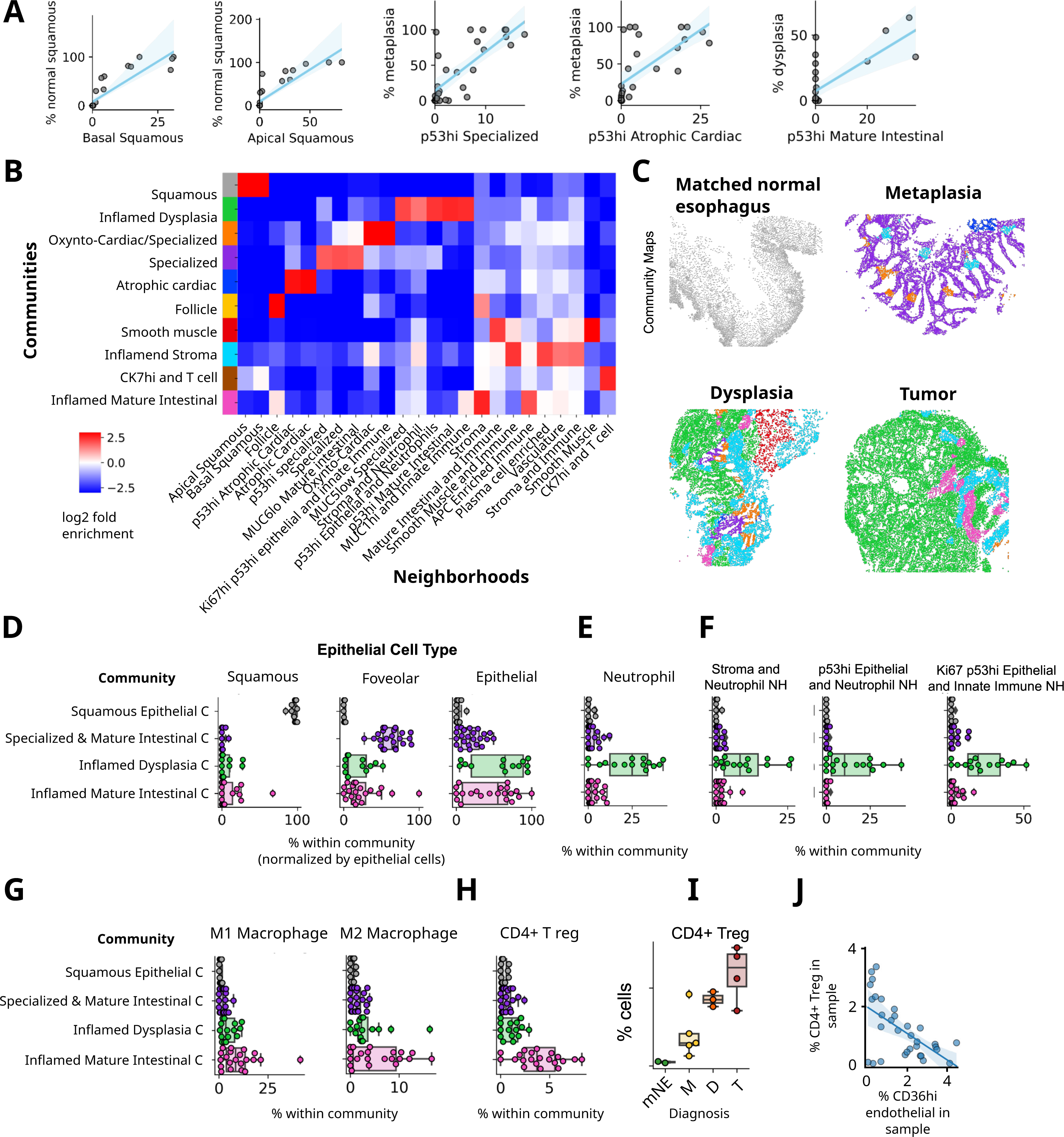
Community (Neighborhood of neighborhoods) analysis of CODEX imaging illustrates the dynamic rearrangements of intricate epithelial-stromal cell entities during BE progression. **(A)** Percentage of epithelial cells as determined by a pathologist (y axis) versus percentage of the neighborhoods as determined by CODEX for each of the 27 regions imaged with correlations. **(B)** Neighborhoods enriched or depleted within individual communities identified. **(C)** Maps of communities mapped back to original coordinates for one representative example donor. Community color code is the same as in B. **(D-E)** Percentage of cell type within each community that correlated with pathologist-classified epithelium for (D) epithelial cell types and (E) neutrophils. **(F)** Percentage of neighborhoods within each community that correlated with pathologist-classified epithelium. **(G-H)** Percentage of cell type within each community that correlated with pathologist-classified epithelium for (G) macrophage phenotypes and (H) CD4+ Tregs. **(I)** Percentage of CD4+ Tregs in progression phases. **(J)** Percentage of CD4+ Tregs versus percentage of CD36hi endothelial cells within the same sample imaged.

**Figure S13.**
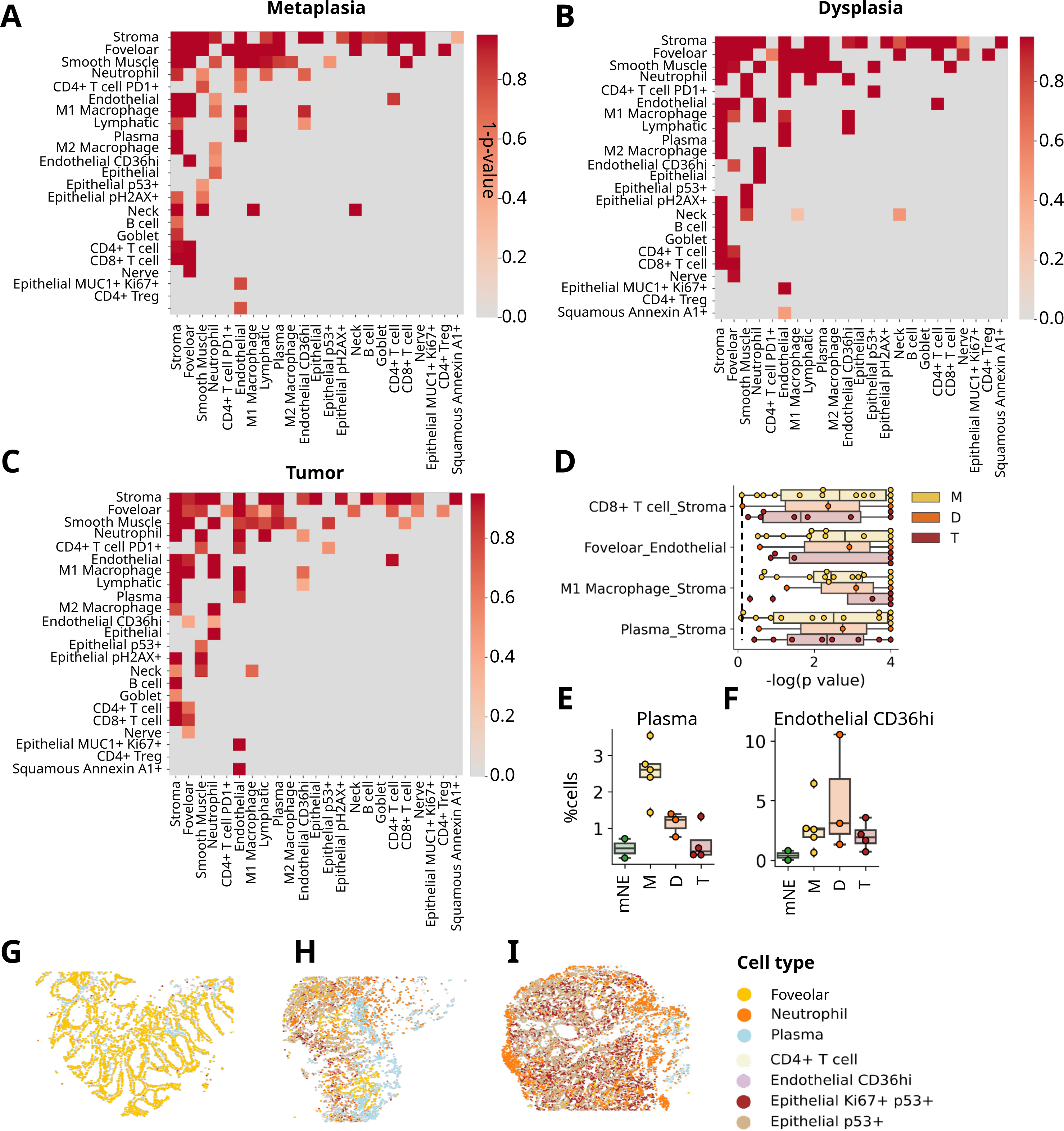
Cell-cell interaction analysis from CODEX images reveals dynamic and increasingly diverse cell interaction pairs during BE progression. **(A-C)** Significant (average *p*-value <0.05) cell-cell interactions shown for (A) metaplasia, (B) dysplasia, and (C) tumor classified samples. **(D)** Cell-cell pairs found enriched in *M*, *D* and *T* disease phases with dashed line representing a p value of 0.05. **(E-F)** Percentage of (E) plasma cells and (F) CD36hi endothelial cells found across different disease phases. **(G-I)** Subset of cells identified by CODEX imaging replotted to tissue coordinates for (G) metaplasia, (H) dysplasia, and (I) tumor samples.

## Supplemental Tables

**Table S1. Patient and biosample metadata.** Patient information, biosample descriptions, and histological diagnosis are provided for 64 samples.

**Table S2. Coarse grained differentially expressed genes.** For each of 16 coarse grained cell types, differential expression was calculated comparing phases of progressing with DESeq2 and the pseudo-bulk transcriptomes per biosample. Results were filtered through association testing with the ambient RNA from empty droplets.

**Table S3. Fine grained gene markers.** Gene markers of fine grained cell types based on statistical differential expression analysis between clusters.

**Table S4. Fibroblast pathway signature scores.** Using the DEGs of fibroblasts, B cells, natural killer cells (NK) and monocyte-derived cells, selected pathways were tested for association using Enrichr. Inputs were DEGs filtered to have at least a log2 fold change of 0.58 and adjusted p-value of 0.05.

**Table S5. Patient cell counts.** From the single cell RNA-seq data, counts of both coarse grain and fine grained cell types per patient.

**Table S6. ECM Proteomic differential abundance statistics.** Comparing ECM protein abundance between pooled Tumor (T) samples versus matched Normal Esophagus (mNE) samples using paired T-tests with multiple testing correction.

**Table S7. ECM Proteomics DIA isolation scheme.** Samples were analyzed for ECM protein abundance by data-independent acquisition (DIA) using variable-sized windows covering the m/z 400-1,250 range.

**Table S8. CODEX Pathology concordance**. Using imaging data from 5 human donors, three pathologists independently evaluated the H&E staining of the sections performed on the same tissue sections that were used for the CODEX multiplexed imaging. Pathologist scores were then aggregated and averaged for disease phase granular diagnosis (e.g., mNE, mNS, M), and estimated percentages of type of epithelium in each image (e.g., % squamous, % metaplasia, % dysplasia, % tumor).

**Table S9. CODEX antibody information.** The antibody panels used in generating CODEX data were chosen to include targets that identify subtypes of intestinal epithelium and stromal cells, and cells of the innate and adaptive immune system.

**Table S10. CODEX communities.** Percentages of cell type composition for each of the 4 communities (mNE: grey; M: purple; D: green; T: pink) that correlated with diseased epithelial, broken down by epithelial, immune, and mesenchymal groupings.

**Table S11. CODEX neighborhood analysis.** Spatial neighborhood (NH) information was used for cell-cell interaction statistics across progression phases (M, D, T). The frequency of neighbors, using a nearest neighbor approach, was compared to the frequency of occurrences in null-models, achieved by 10,000 permutations of cell type locations.

## REFERENCES

1. Uhlenhopp, D.J., Then, E.O., Sunkara, T., and Gaduputi, V. (2020). Epidemiology of esophageal cancer: update in global trends, etiology and risk factors. Clin. J. Gastroenterol. 13, 1010–1021. 10.1007/s12328-020-01237-x.

2. Smyth, E.C., Lagergren, J., Fitzgerald, R.C., Lordick, F., Shah, M.A., Lagergren, P., and Cunningham, D. (2017). Oesophageal cancer. Nat. Rev. Dis. Primer 3, 17048. 10.1038/nrdp.2017.48.

3. Yang, J., Liu, X., Cao, S., Dong, X., Rao, S., and Cai, K. (2020). Understanding Esophageal Cancer: The Challenges and Opportunities for the Next Decade. Front. Oncol. 10, 1727. 10.3389/fonc.2020.01727.

4. McColl, K.E.L. (2019). What is causing the rising incidence of esophageal adenocarcinoma in the West and will it also happen in the East? J. Gastroenterol. 54, 669–673. 10.1007/s00535-019-01593-7.

5. Hur, C., Miller, M., Kong, C.Y., Dowling, E.C., Nattinger, K.J., Dunn, M., and Feuer, E.J. (2013). Trends in Esophageal Adenocarcinoma Incidence and Mortality. Cancer 119, 1149–1158. 10.1002/cncr.27834.

6.. Cancer Genome Atlas Research Network, Analysis Working Group: Asan University, BC Cancer Agency, Brigham and Women’s Hospital, Broad Institute, Brown University, Case Western Reserve University, Dana-Farber Cancer Institute, Duke University, Greater Poland Cancer Centre, et al. (2017). Integrated genomic characterization of oesophageal carcinoma. Nature 541, 169–175. 10.1038/nature20805.

7. Nowicki-Osuch, K., Zhuang, L., Jammula, S., Bleaney, C.W., Mahbubani, K.T., Devonshire, G., Katz-Summercorn, A., Eling, N., Wilbrey-Clark, A., Madissoon, E., et al. (2021). Molecular phenotyping reveals the identity of Barrett’s esophagus and its malignant transition. Science 373, 760–767. 10.1126/science.abd1449.

8. Krishnamoorthi, R., Mohan, B.P., Jayaraj, M., Wang, K.K., Katzka, D.A., Ross, A., Adler, D.G., and Iyer, P.G. (2020). Risk of progression in Barrett’s esophagus indefinite for dysplasia: a systematic review and meta-analysis. Gastrointest. Endosc. 91, 3–10.e3. 10.1016/j.gie.2019.07.037.

9. Weaver, J.M.J., Ross-Innes, C.S., Shannon, N., Lynch, A.G., Forshew, T., Barbera, M., Murtaza, M., Ong, C.-A.J., Lao-Sirieix, P., Dunning, M.J., et al. (2014). Ordering of mutations in preinvasive disease stages of esophageal carcinogenesis. Nat. Genet. 46, 837–843. 10.1038/ng.3013.

10. Killcoyne, S., Gregson, E., Wedge, D.C., Woodcock, D.J., Eldridge, M.D., de la Rue, R., Miremadi, A., Abbas, S., Blasko, A., Kosmidou, C., et al. (2020). Genomic copy number predicts esophageal cancer years before transformation. Nat. Med. 26, 1726–1732. 10.1038/s41591-020-1033-y.

11. Katz-Summercorn, A.C., Jammula, S., Frangou, A., Peneva, I., O’Donovan, M., Tripathi, M., Malhotra, S., di Pietro, M., Abbas, S., Devonshire, G., et al. (2022). Multi-omic cross-sectional cohort study of pre-malignant Barrett’s esophagus reveals early structural variation and retrotransposon activity. Nat. Commun. 13, 1407. 10.1038/s41467-022-28237-4.

12. Stachler, M.D., Camarda, N.D., Deitrick, C., Kim, A., Agoston, A.T., Odze, R.D., Hornick, J.L., Nag, A., Thorner, A.R., Ducar, M., et al. (2018). Detection of Mutations in Barrett’s Esophagus Before Progression to High-Grade Dysplasia or Adenocarcinoma. Gastroenterology 155, 156–167. 10.1053/j.gastro.2018.03.047.

13. Luebeck, J., Ng, A.W.T., Galipeau, P.C., Li, X., Sanchez, C.A., Katz-Summercorn, A.C., Kim, H., Jammula, S., He, Y., Lippman, S.M., et al. (2023). Extrachromosomal DNA in the cancerous transformation of Barrett’s oesophagus. Nature. 10.1038/s41586-023-05937-5.

14. Maag, J.L.V., Fisher, O.M., Levert-Mignon, A., Kaczorowski, D.C., Thomas, M.L., Hussey, D.J., Watson, D.I., Wettstein, A., Bobryshev, Y.V., Edwards, M., et al. (2017). Novel Aberrations Uncovered in Barrett’s Esophagus and Esophageal Adenocarcinoma Using Whole Transcriptome Sequencing. Mol. Cancer Res. MCR 15, 1558–1569. 10.1158/1541-7786.MCR-17-0332.

15. Nowicki-Osuch, K., Zhuang, L., Cheung, T.S., Black, E.L., Masque-Soler, N., Devonshire, G., Redmond, A.M., Freeman, A., di Pietro, M., Pilonis, N., et al. (2023). Single-cell RNA sequencing unifies developmental programs of Esophageal and Gastric Intestinal Metaplasia. Cancer Discov., CD-22–0824. 10.1158/2159-8290.CD-22-0824.

16. Murphy, K.J., Chambers, C.R., Herrmann, D., Timpson, P., and Pereira, B.A. (2021). Dynamic Stromal Alterations Influence Tumor-Stroma Crosstalk to Promote Pancreatic Cancer and Treatment Resistance. Cancers 13, 3481. 10.3390/cancers13143481.

17. Mishra, R., Haldar, S., Placencio, V., Madhav, A., Rohena-Rivera, K., Agarwal, P., Duong, F., Angara, B., Tripathi, M., Liu, Z., et al. Stromal epigenetic alterations drive metabolic and neuroendocrine prostate cancer reprogramming. J. Clin. Invest. 128, 4472–4484. 10.1172/JCI99397.

18. Xu, M., Zhang, T., Xia, R., Wei, Y., and Wei, X. (2022). Targeting the tumor stroma for cancer therapy. Mol. Cancer 21, 208. 10.1186/s12943-022-01670-1.

19. Greten, F.R., and Grivennikov, S.I. (2019). Inflammation and Cancer: Triggers, Mechanisms, and Consequences. Immunity 51, 27–41. 10.1016/j.immuni.2019.06.025.

20. Korsunsky, I., Millard, N., Fan, J., Slowikowski, K., Zhang, F., Wei, K., Baglaenko, Y., Brenner, M., Loh, P.-R., and Raychaudhuri, S. (2019). Fast, sensitive and accurate integration of single-cell data with Harmony. Nat. Methods 16, 1289–1296. 10.1038/s41592-019-0619-0.

21. Han, X., Zhou, Z., Fei, L., Sun, H., Wang, R., Chen, Y., Chen, H., Wang, J., Tang, H., Ge, W., et al. (2020). Construction of a human cell landscape at single-cell level. Nature 581, 303–309. 10.1038/s41586-020-2157-4.

22. Parikh, K., Antanaviciute, A., Fawkner-Corbett, D., Jagielowicz, M., Aulicino, A., Lagerholm, C., Davis, S., Kinchen, J., Chen, H.H., Alham, N.K., et al. (2019). Colonic epithelial cell diversity in health and inflammatory bowel disease. Nature 567, 49–55. 10.1038/s41586-019-0992-y.

23. Gutierrez-Gonzalez, L., Graham, T.A., Rodriguez-Justo, M., Leedham, S.J., Novelli, M.R., Gay, L.J., Ventayol-Garcia, T., Green, A., Mitchell, I., Stoker, D.L., et al. (2011). The clonal origins of dysplasia from intestinal metaplasia in the human stomach. Gastroenterology 140, 1251–1260.e1-6. 10.1053/j.gastro.2010.12.051.

24. Squair, J.W., Gautier, M., Kathe, C., Anderson, M.A., James, N.D., Hutson, T.H., Hudelle, R., Qaiser, T., Matson, K.J.E., Barraud, Q., et al. (2021). Confronting false discoveries in single-cell differential expression. Nat. Commun. 12, 5692. 10.1038/s41467-021-25960-2.

25. Love, M.I., Huber, W., and Anders, S. (2014). Moderated estimation of fold change and dispersion for RNA-seq data with DESeq2. Genome Biol. 15, 550. 10.1186/s13059-014-0550-8.

26. Xie, Z., Bailey, A., Kuleshov, M.V., Clarke, D.J.B., Evangelista, J.E., Jenkins, S.L., Lachmann, A., Wojciechowicz, M.L., Kropiwnicki, E., Jagodnik, K.M., et al. (2021). Gene Set Knowledge Discovery with Enrichr. Curr. Protoc. 1, e90. 10.1002/cpz1.90.

27. Chen, E.Y., Tan, C.M., Kou, Y., Duan, Q., Wang, Z., Meirelles, G.V., Clark, N.R., and Ma’ayan, A. (2013). Enrichr: interactive and collaborative HTML5 gene list enrichment analysis tool. BMC Bioinformatics 14, 128. 10.1186/1471-2105-14-128.

28. McDonald, S.A.C., Lavery, D., Wright, N.A., and Jansen, M. (2015). Barrett oesophagus: lessons on its origins from the lesion itself. Nat. Rev. Gastroenterol. Hepatol. 12, 50–60. 10.1038/nrgastro.2014.181.

29. Barbera, M., and Fitzgerald, R.C. (2010). Cellular origin of Barrett’s metaplasia and oesophageal stem cells. Biochem. Soc. Trans. 38, 370–373. 10.1042/BST0380370.

30. Jiang, M., Li, H., Zhang, Y., Yang, Y., Lu, R., Liu, K., Lin, S., Lan, X., Wang, H., Wu, H., et al. (2017). Transitional basal cells at the squamous–columnar junction generate Barrett’s oesophagus. Nature 550, 529–533. 10.1038/nature24269.

31. Hanahan, D. (2022). Hallmarks of Cancer: New Dimensions. Cancer Discov. 12, 31–46. 10.1158/2159-8290.CD-21-1059.

32. Buechler, M.B., Pradhan, R.N., Krishnamurty, A.T., Cox, C., Calviello, A.K., Wang, A.W., Yang, Y.A., Tam, L., Caothien, R., Roose-Girma, M., et al. (2021). Cross-tissue organization of the fibroblast lineage. Nature 593, 575–579. 10.1038/s41586-021-03549-5.

33. Lendahl, U., Muhl, L., and Betsholtz, C. (2022). Identification, discrimination and heterogeneity of fibroblasts. Nat. Commun. 13, 3409. 10.1038/s41467-022-30633-9.

34. Lee, S.-H., Contreras Panta, E.W., Gibbs, D., Won, Y., Min, J., Zhang, C., Roland, J.T., Hong, S.-H., Sohn, Y., Krystofiak, E., et al. (2023). Apposition of fibroblasts with metaplastic gastric cells promotes dysplastic transition. Gastroenterology, S001650852300731X. 10.1053/j.gastro.2023.04.038.

35. De Palma, M., Biziato, D., and Petrova, T.V. (2017). Microenvironmental regulation of tumour angiogenesis. Nat. Rev. Cancer 17, 457–474. 10.1038/nrc.2017.51.

36. Raza, A., Franklin, M.J., and Dudek, A.Z. (2010). Pericytes and vessel maturation during tumor angiogenesis and metastasis. Am. J. Hematol. 85, 593–598. 10.1002/ajh.21745.

37. Dias, D.O., Kalkitsas, J., Kelahmetoglu, Y., Estrada, C.P., Tatarishvili, J., Holl, D., Jansson, L., Banitalebi, S., Amiry-Moghaddam, M., Ernst, A., et al. (2021). Pericyte-derived fibrotic scarring is conserved across diverse central nervous system lesions. Nat. Commun. 12, 5501. 10.1038/s41467-021-25585-5.

38. He, S., Wang, L.-H., Liu, Y., Li, Y.-Q., Chen, H.-T., Xu, J.-H., Peng, W., Lin, G.-W., Wei, P.-P., Li, B., et al. (2020). Single-cell transcriptome profiling of an adult human cell atlas of 15 major organs. Genome Biol. 21, 294. 10.1186/s13059-020-02210-0.

39. Bosma, E.K., van Noorden, C.J.F., Schlingemann, R.O., and Klaassen, I. (2018). The role of plasmalemma vesicle-associated protein in pathological breakdown of blood-brain and blood-retinal barriers: potential novel therapeutic target for cerebral edema and diabetic macular edema. Fluids Barriers CNS 15, 24. 10.1186/s12987-018-0109-2.

40. Schupp, J.C., Adams, T.S., Cosme, C., Raredon, M.S.B., Yuan, Y., Omote, N., Poli, S., Chioccioli, M., Rose, K.-A., Manning, E.P., et al. (2021). Integrated Single-Cell Atlas of Endothelial Cells of the Human Lung. Circulation 144, 286–302. 10.1161/CIRCULATIONAHA.120.052318.

41. Lee, H.-W., Xu, Y., He, L., Choi, W., Gonzalez, D., Jin, S.-W., and Simons, M. (2021). Role of Venous Endothelial Cells in Developmental and Pathologic Angiogenesis. Circulation 144, 1308–1322. 10.1161/CIRCULATIONAHA.121.054071.

42. Leu, A.J., Berk, D.A., Lymboussaki, A., Alitalo, K., and Jain, R.K. (2000). Absence of functional lymphatics within a murine sarcoma: a molecular and functional evaluation. Cancer Res. 60, 4324–4327.

43. Padera, T.P., Kadambi, A., di Tomaso, E., Carreira, C.M., Brown, E.B., Boucher, Y., Choi, N.C., Mathisen, D., Wain, J., Mark, E.J., et al. (2002). Lymphatic metastasis in the absence of functional intratumor lymphatics. Science 296, 1883–1886. 10.1126/science.1071420.

44. Crome, S.Q., Nguyen, L.T., Lopez-Verges, S., Yang, S.Y.C., Martin, B., Yam, J.Y., Johnson, D.J., Nie, J., Pniak, M., Yen, P.H., et al. (2017). A distinct innate lymphoid cell population regulates tumor-associated T cells. Nat. Med. 23, 368–375. 10.1038/nm.4278.

45. Corvino, D., Kumar, A., and Bald, T. (2022). Plasticity of NK cells in Cancer. Front. Immunol. 13, 888313. 10.3389/fimmu.2022.888313.

46. Fristedt, R., Borg, D., Hedner, C., Berntsson, J., Nodin, B., Eberhard, J., Micke, P., and Jirström, K. (2016). Prognostic impact of tumour-associated B cells and plasma cells in oesophageal and gastric adenocarcinoma. J. Gastrointest. Oncol. 7, 848–859. 10.21037/jgo.2016.11.07.

47. Li, X., Paulson, T.G., Galipeau, P.C., Sanchez, C.A., Liu, K., Kuhner, M.K., Maley, C.C., Self, S.G., Vaughan, T.L., Reid, B.J., et al. (2015). Assessment of Esophageal Adenocarcinoma Risk Using Somatic Chromosome Alterations in Longitudinal Samples in Barrett’s Esophagus. Cancer Prev. Res. Phila. Pa 8, 845–856. 10.1158/1940-6207.CAPR-15-0130.

48. Tickle T, Tirosh I, Georgescu C, Brown M, Haas B (2019). inferCNV of the Trinity CTAT Project.

49. Paulson, T.G., Maley, C.C., Li, X., Li, H., Sanchez, C.A., Chao, D.L., Odze, R.D., Vaughan, T.L., Blount, P.L., and Reid, B.J. (2009). Chromosomal instability and copy number alterations in Barrett’s esophagus and esophageal adenocarcinoma. Clin. Cancer Res. Off. J. Am. Assoc. Cancer Res. 15, 3305–3314. 10.1158/1078-0432.CCR-08-2494.

50. Stachler, M.D., Taylor-Weiner, A., Peng, S., McKenna, A., Agoston, A.T., Odze, R.D., Davison, J.M., Nason, K.S., Loda, M., Leshchiner, I., et al. (2015). Paired exome analysis of Barrett’s esophagus and adenocarcinoma. Nat. Genet. 47, 1047–1055. 10.1038/ng.3343.

51. Shimshoni, E., Merry, G.E., Milot, Z.D., Oh, C.Y., Horvath, V., Gould, R.A., Caruso, J.A., Chen-Tanyolac, C., Gascard, P., Sangwan, V., et al. (2023). Epithelial-Stromal Interactions in Barrett’s Esophagus Modeled in Human Organ Chips. Gastro Hep Adv., S2772572323000444. 10.1016/j.gastha.2023.03.009.

52. Norreen-Thorsen, M., Struck, E.C., Öling, S., Zwahlen, M., Von Feilitzen, K., Odeberg, J., Lindskog, C., Pontén, F., Uhlén, M., Dusart, P.J., et al. (2022). A human adipose tissue cell-type transcriptome atlas. Cell Rep. 40, 111046. 10.1016/j.celrep.2022.111046.

53. Wynn, T.A. (2008). Cellular and molecular mechanisms of fibrosis. J. Pathol. 214, 199– 210. 10.1002/path.2277.

54. Wells, R.G. (2008). Cellular sources of extracellular matrix in hepatic fibrosis. Clin. Liver Dis. 12, 759–768, viii. 10.1016/j.cld.2008.07.008.

55. Saadi, A., Das, M., Clemons, N., Zhang, C., Ferguson, M., Tokiwa, G., Serikawa, K., Hardwick, J., Dai, H., Carlini, L., et al. (2014). A potential role for Periostin in Barrett’s carcinogenesis. Cancer Epidemiol. Biomarkers Prev. 16, B28.

56. Onieva, J.L., Xiao, Q., Berciano-Guerrero, M.-Á., Laborda-Illanes, A., de Andrea, C., Chaves, P., Piñeiro, P., Garrido-Aranda, A., Gallego, E., Sojo, B., et al. (2022). High IGKC-Expressing Intratumoral Plasma Cells Predict Response to Immune Checkpoint Blockade. Int. J. Mol. Sci. 23, 9124. 10.3390/ijms23169124.

57. Schmidt, M., Micke, P., Gehrmann, M., and Hengstler, J.G. (2012). Immunoglobulin kappa chain as an immunologic biomarker of prognosis and chemotherapy response in solid tumors. Oncoimmunology 1, 1156–1158. 10.4161/onci.21653.

58. Fiore, V.F., Krajnc, M., Quiroz, F.G., Levorse, J., Pasolli, H.A., Shvartsman, S.Y., and Fuchs, E. (2020). Mechanics of a multilayer epithelium instruct tumour architecture and function. Nature 585, 433–439. 10.1038/s41586-020-2695-9.

59. Cross, S.E., Jin, Y.-S., Rao, J., and Gimzewski, J.K. (2007). Nanomechanical analysis of cells from cancer patients. Nat. Nanotechnol. 2, 780–783. 10.1038/nnano.2007.388.

60. Iyer, S., Gaikwad, R.M., Subba-Rao, V., Woodworth, C.D., and Sokolov, I. (2009). Atomic force microscopy detects differences in the surface brush of normal and cancerous cells. Nat. Nanotechnol. 4, 389–393. 10.1038/nnano.2009.77.

61. Butcher, D.T., Alliston, T., and Weaver, V.M. (2009). A tense situation: forcing tumour progression. Nat. Rev. Cancer 9, 108–122. 10.1038/nrc2544.

62. Kennedy-Darling, J., Bhate, S.S., Hickey, J.W., Black, S., Barlow, G.L., Vazquez, G., Venkataraaman, V.G., Samusik, N., Goltsev, Y., Schürch, C.M., et al. (2021). Highly multiplexed tissue imaging using repeated oligonucleotide exchange reaction. Eur. J. Immunol. 51, 1262–1277. 10.1002/eji.202048891.

63. Black, S., Phillips, D., Hickey, J.W., Kennedy-Darling, J., Venkataraaman, V.G., Samusik, N., Goltsev, Y., Schürch, C.M., and Nolan, G.P. (2021). CODEX multiplexed tissue imaging with DNA-conjugated antibodies. Nat. Protoc. 16, 3802–3835. 10.1038/s41596-021-00556-8.

64. Hickey, J.W., Neumann, E.K., Radtke, A.J., Camarillo, J.M., Beuschel, R.T., Albanese, A., McDonough, E., Hatler, J., Wiblin, A.E., Fisher, J., et al. (2022). Spatial mapping of protein composition and tissue organization: a primer for multiplexed antibody-based imaging. Nat. Methods 19, 284–295. 10.1038/s41592-021-01316-y.

65. Hickey, J.W., Becker, W.R., Nevins, S.A., Horning, A., Perez, A.E., Chiu, R., Chen, D.C., Cotter, D., Esplin, E.D., Weimer, A.K., et al. (2021). 35672395. 2021.11.25.469203. 10.1101/2021.11.25.469203.

66. Souza, R.F., and Spechler, S.J. (2022). Mechanisms and pathophysiology of Barrett oesophagus. Nat. Rev. Gastroenterol. Hepatol. 19, 605–620. 10.1038/s41575-022-00622- w.

67. Hickey, J.W., Tan, Y., Nolan, G.P., and Goltsev, Y. (2021). Strategies for Accurate Cell Type Identification in CODEX Multiplexed Imaging Data. Front. Immunol. 12, 727626. 10.3389/fimmu.2021.727626.

68. D’Alessio, F.R., Tsushima, K., Aggarwal, N.R., West, E.E., Willett, M.H., Britos, M.F., Pipeling, M.R., Brower, R.G., Tuder, R.M., McDyer, J.F., et al. (2009). CD4+CD25+Foxp3+ Tregs resolve experimental lung injury in mice and are present in humans with acute lung injury. J. Clin. Invest. 119, 2898–2913. 10.1172/JCI36498.

69. DeFilippis, R.A., Chang, H., Dumont, N., Rabban, J.T., Chen, Y.-Y., Fontenay, G.V., Berman, H.K., Gauthier, M.L., Zhao, J., Hu, D., et al. (2012). CD36 repression activates a multicellular stromal program shared by high mammographic density and tumor tissues. Cancer Discov. 2, 826–839. 10.1158/2159-8290.CD-12-0107.

70. McDonald, S.A.C., Graham, T.A., Lavery, D.L., Wright, N.A., and Jansen, M. (2015). The Barrett’s Gland in Phenotype Space. Cell. Mol. Gastroenterol. Hepatol. 1, 41–54. 10.1016/j.jcmgh.2014.10.001.

71. Calon, A., Tauriello, D.V.F., and Batlle, E. (2014). TGF-beta in CAF-mediated tumor growth and metastasis. Semin. Cancer Biol. 25, 15–22. 10.1016/j.semcancer.2013.12.008.

72. DeFilippis, R.A., Fordyce, C., Patten, K., Chang, H., Zhao, J., Fontenay, G.V., Kerlikowske, K., Parvin, B., and Tlsty, T.D. (2014). Stress signaling from human mammary epithelial cells contributes to phenotypes of mammographic density. Cancer Res. 74, 5032–5044. 10.1158/0008-5472.CAN-13-3390.

73. Hao, Y., Triadafilopoulos, G., Sahbaie, P., Young, H.S., Omary, M.B., and Lowe, A.W. (2006). Gene expression profiling reveals stromal genes expressed in common between Barrett’s esophagus and adenocarcinoma. Gastroenterology 131, 925–933. 10.1053/j.gastro.2006.04.026.

74. Tilman, G., Mattiussi, M., Brasseur, F., van Baren, N., and Decottignies, A. (2007). Human periostin gene expression in normal tissues, tumors and melanoma: evidences for periostin production by both stromal and melanoma cells. Mol. Cancer 6, 80. 10.1186/1476-4598-6-80.

75. Park, J., and Scherer, P.E. (2012). Endotrophin - a novel factor linking obesity with aggressive tumor growth. Oncotarget 3, 1487–1488. 10.18632/oncotarget.796.

76. Williams, L., Layton, T., Yang, N., Feldmann, M., and Nanchahal, J. (2022). Collagen VI as a driver and disease biomarker in human fibrosis. FEBS J. 289, 3603–3629. 10.1111/febs.16039.

77. Davidson, S., Efremova, M., Riedel, A., Mahata, B., Pramanik, J., Huuhtanen, J., Kar, G., Vento-Tormo, R., Hagai, T., Chen, X., et al. (2020). Single-Cell RNA Sequencing Reveals a Dynamic Stromal Niche That Supports Tumor Growth. Cell Rep. 31, 107628. 10.1016/j.celrep.2020.107628.

78. Fuhrmann, A., Staunton, J.R., Nandakumar, V., Banyai, N., Davies, P.C.W., and Ros, R. (2011). AFM stiffness nanotomography of normal, metaplastic and dysplastic human esophageal cells. Phys. Biol. 8, 015007. 10.1088/1478-3975/8/1/015007.

79. Northcott, J.M., Dean, I.S., Mouw, J.K., and Weaver, V.M. (2018). Feeling Stress: The Mechanics of Cancer Progression and Aggression. Front. Cell Dev. Biol. 6, 17. 10.3389/fcell.2018.00017.

80. Li, J., Tan, J., Martino, M.M., and Lui, K.O. (2018). Regulatory T-Cells: Potential Regulator of Tissue Repair and Regeneration. Front. Immunol. 9, 585. 10.3389/fimmu.2018.00585.

81. Lagisetty, K.H., McEwen, D.P., Nancarrow, D.J., Schiebel, J.G., Ferrer-Torres, D., Ray, D., Frankel, T.L., Lin, J., Chang, A.C., Kresty, L.A., et al. (2021). Immune determinants of Barrett’s progression to esophageal adenocarcinoma. JCI Insight 6, e143888, 143888. 10.1172/jci.insight.143888.

82. Gokon, Y., Fujishima, F., Taniyama, Y., Ishida, H., Yamagata, T., Sawai, T., Uzuki, M., Ichikawa, H., Itakura, Y., Takahashi, K., et al. (2020). Immune microenvironment in Barrett’s esophagus adjacent to esophageal adenocarcinoma: possible influence of adjacent mucosa on cancer development and progression. Virchows Arch. Int. J. Pathol. 477, 825–834. 10.1007/s00428-020-02854-0.

83. Wang, Z., Xiong, S., Mao, Y., Chen, M., Ma, X., Zhou, X., Ma, Z., Liu, F., Huang, Z., Luo, Q., et al. (2016). Periostin promotes immunosuppressive premetastatic niche formation to facilitate breast tumour metastasis. J. Pathol. 239, 484–495. 10.1002/path.4747.

84. Gowhari Shabgah, A., Haleem Al-Qaim, Z., Markov, A., Valerievich Yumashev, A., Ezzatifar, F., Ahmadi, M., Mohammad Gheibihayat, S., and Gholizadeh Navashenaq, J. (2021). Chemokine CXCL14; a double-edged sword in cancer development. Int. Immunopharmacol. 97, 107681. 10.1016/j.intimp.2021.107681.

85. Morra, L., and Moch, H. (2011). Periostin expression and epithelial-mesenchymal transition in cancer: a review and an update. Virchows Arch. Int. J. Pathol. 459, 465–475. 10.1007/s00428-011-1151-5.

86. Martin, M. Cutadapt Removes Adapter Sequences From High-Throughput Sequencing Reads. EMBnet.journal 17, 10–12. 10.14806/ej.17.1.200.

87. Bray, N.L., Pimentel, H., Melsted, P., and Pachter, L. (2016). Near-optimal probabilistic RNA-seq quantification. Nat. Biotechnol. 34, 525–527. 10.1038/nbt.3519.

88. Melsted, P., Booeshaghi, A.S., Liu, L., Gao, F., Lu, L., Min, K.H., da Veiga Beltrame, E., Hjörleifsson, K.E., Gehring, J., and Pachter, L. (2021). Modular, efficient and constant-memory single-cell RNA-seq preprocessing. Nat. Biotechnol. 39, 813–818. 10.1038/s41587-021-00870-2.

89. Wolf, F.A., Angerer, P., and Theis, F.J. (2018). SCANPY: large-scale single-cell gene expression data analysis. Genome Biol. 19, 15. 10.1186/s13059-017-1382-0.

90. Wolock, S.L., Lopez, R., and Klein, A.M. (2019). Scrublet: Computational Identification of Cell Doublets in Single-Cell Transcriptomic Data. Cell Syst. 8, 281–291.e9. 10.1016/j.cels.2018.11.005.

91. Zheng, G.X.Y., Terry, J.M., Belgrader, P., Ryvkin, P., Bent, Z.W., Wilson, R., Ziraldo, S.B., Wheeler, T.D., McDermott, G.P., Zhu, J., et al. (2017). Massively parallel digital transcriptional profiling of single cells. Nat. Commun. 8, 14049. 10.1038/ncomms14049.

92. McInnes, L., Healy, J., and Melville, J. (2018). UMAP: Uniform Manifold Approximation and Projection for Dimension Reduction. ArXiv180203426 Cs Stat.

93. Wolf, F.A., Hamey, F.K., Plass, M., Solana, J., Dahlin, J.S., Göttgens, B., Rajewsky, N., Simon, L., and Theis, F.J. (2019). PAGA: graph abstraction reconciles clustering with trajectory inference through a topology preserving map of single cells. Genome Biol. 20, 59. 10.1186/s13059-019-1663-x.

94. Traag, V.A., Waltman, L., and van Eck, N.J. (2019). From Louvain to Leiden: guaranteeing well-connected communities. Sci. Rep. 9, 5233. 10.1038/s41598-019-41695-z.

95. Speir, M.L., Bhaduri, A., Markov, N.S., Moreno, P., Nowakowski, T.J., Papatheodorou, I., Pollen, A.A., Raney, B.J., Seninge, L., Kent, W.J., et al. (2021). UCSC Cell Browser: visualize your single-cell data. Bioinformatics 37, 4578–4580. 10.1093/bioinformatics/btab503.

96. Osumi-Sutherland, D., Xu, C., Keays, M., Levine, A.P., Kharchenko, P.V., Regev, A., Lein, E., and Teichmann, S.A. (2021). Cell type ontologies of the Human Cell Atlas. Nat. Cell Biol. 23, 1129–1135. 10.1038/s41556-021-00787-7.

97. Madissoon, E., Wilbrey-Clark, A., Miragaia, R.J., Saeb-Parsy, K., Mahbubani, K.T., Georgakopoulos, N., Harding, P., Polanski, K., Huang, N., Nowicki-Osuch, K., et al. (2019). scRNA-seq assessment of the human lung, spleen, and esophagus tissue stability after cold preservation. Genome Biol. 21, 1. 10.1186/s13059-019-1906-x.

98. Yang, S., Corbett, S.E., Koga, Y., Wang, Z., Johnson, W.E., Yajima, M., and Campbell, J.D. (2020). Decontamination of ambient RNA in single-cell RNA-seq with DecontX. Genome Biol. 21, 57. 10.1186/s13059-020-1950-6.

99. Büttner, M., Ostner, J., Müller, C.L., Theis, F.J., and Schubert, B. (2021). scCODA is a Bayesian model for compositional single-cell data analysis. Nat. Commun. 12, 6876. 10.1038/s41467-021-27150-6.

100. Sturm, G. (2022). infercnvpy: Scanpy plugin to infer copy number variation (CNV) from single-cell transcriptomics data.

101. Data-independent acquisition and quantification of extracellular matrix from human lung in chronic inflammation-associated carcinomas - PubMed https://pubmed.ncbi.nlm.nih.gov/36228107/.

102. Escher, C., Reiter, L., MacLean, B., Ossola, R., Herzog, F., Chilton, J., MacCoss, M.J., and Rinner, O. (2012). Using iRT, a normalized retention time for more targeted measurement of peptides. Proteomics 12, 1111–1121. 10.1002/pmic.201100463.

103. Gillet, L.C., Navarro, P., Tate, S., Röst, H., Selevsek, N., Reiter, L., Bonner, R., and Aebersold, R. (2012). Targeted data extraction of the MS/MS spectra generated by data-independent acquisition: a new concept for consistent and accurate proteome analysis. Mol. Cell. Proteomics MCP 11, O111.016717. 10.1074/mcp.O111.016717.

104. Collins, B.C., Hunter, C.L., Liu, Y., Schilling, B., Rosenberger, G., Bader, S.L., Chan, D.W., Gibson, B.W., Gingras, A.-C., Held, J.M., et al. (2017). Multi-laboratory assessment of reproducibility, qualitative and quantitative performance of SWATH-mass spectrometry. Nat. Commun. 8, 291. 10.1038/s41467-017-00249-5.

105. Schilling, B., Gibson, B.W., and Hunter, C.L. (2017). Generation of High-Quality SWATH® Acquisition Data for Label-free Quantitative Proteomics Studies Using TripleTOF® Mass Spectrometers. Methods Mol. Biol. Clifton NJ 1550, 223–233. 10.1007/978-1-4939-6747-6_16.

106. Rosenberger, G., Koh, C.C., Guo, T., Röst, H.L., Kouvonen, P., Collins, B.C., Heusel, M., Liu, Y., Caron, E., Vichalkovski, A., et al. (2014). A repository of assays to quantify 10,000 human proteins by SWATH-MS. Sci. Data 1, 140031. 10.1038/sdata.2014.31.

107. Storey, J.D. (2002). A direct approach to false discovery rates. J. R. Stat. Soc. Ser. B Stat. Methodol. 64, 479–498.

108. Lee, M.Y., Bedia, J.S., Bhate, S.S., Barlow, G.L., Phillips, D., Fantl, W.J., Nolan, G.P., and Schürch, C.M. (2022). CellSeg: a robust, pre-trained nucleus segmentation and pixel quantification software for highly multiplexed fluorescence images. BMC Bioinformatics 23, 46. 10.1186/s12859-022-04570-9.

109. Bhate, S.S., Barlow, G.L., Schürch, C.M., and Nolan, G.P. (2022). Tissue schematics map the specialization of immune tissue motifs and their appropriation by tumors. Cell Syst. 13, 109–130.e6. 10.1016/j.cels.2021.09.012.

110. Schürch, C.M., Bhate, S.S., Barlow, G.L., Phillips, D.J., Noti, L., Zlobec, I., Chu, P., Black, S., Demeter, J., McIlwain, D.R., et al. (2020). Coordinated Cellular Neighborhoods Orchestrate Antitumoral Immunity at the Colorectal Cancer Invasive Front. Cell 182, 1341–1359.e19. 10.1016/j.cell.2020.07.005.

